# The dynamics of adaptation to stress from standing genetic variation and *de novo* mutations

**DOI:** 10.1101/2022.03.26.485920

**Authors:** S. Lorena Ament-Velásquez, Ciaran Gilchrist, Alexandre Rêgo, Devin P Bendixsen, Claire Brice, Julie Michelle Grosse-Sommer, Nima Rafati, Rike Stelkens

## Abstract

Adaptation from standing genetic variation is an important process underlying evolution in natural populations but we rarely get the opportunity to observe the dynamics of fitness changes in real time. Here, we used the power of microbial experimental evolution and whole population sequencing to track the phenotypic and genomic changes of genetically diverse yeast populations in environments with different stress levels. We found that populations rapidly and in parallel increased in fitness in stressful environments. The founder’s genetic diversity was quickly depleted, however, not to the same degree in all populations and environments. Some populations fixed all ancestral variation in < 30 generations while others maintained diversity across hundreds of generations. We also observed parallelism at the gene and pathway level. Specifically, we detected up to seven genes harbouring multiple independent mutations in different populations, and a general enrichment for mutations affecting downstream effectors of the high-osmolarity-glycerol pathway in three out of four environments. Adaptation to the most stressful environment was characterised by the fast evolution of functional haploidy, likely driven by standing genetic variation. Almost 40% of all populations contained aneuploidies (losses or gains of chromosomes) at least once during experimental evolution. Some aneuploidies were maintained for hundreds of generations in parallel in different replicates, suggesting they were adaptive. This work shows that experimental evolution is a great tool to address the interplay between standing variation and the influx of *de novo* mutations, leading to a better understanding of the demographic and environmental drivers and constraints of a population’s capacity to adapt to environmental change.

## Background

Ongoing climate change drives species outside of their ecological comfort zone and rates of environmental change often exceed rates of adaptation (Radchuk et al., 2019). To better manage at-risk populations, we need to understand adaptation dynamics from different sources of variation, and under different types of selection pressures (Bitter et al., 2019; Chaturvedi et al., 2021; Lai et al., 2019; Matuszewski et al., 2015). Experimental evolution with microbial model systems paired with time-series whole genome sequencing is a powerful tool to study adaptation and the underlying genetic mechanisms leading to fitness change. Most experiments start from clonal populations, i.e. initially isogenic lineages adapting and diverging over time by accumulating new mutations (e.g. Good et al., 2017; Johnson et al., 2021; Lenski et al., 1991). This allows for the modelling of adaptation dynamics by selection on *de novo* mutations as is the case in highly homogenous populations, e.g. clonal pathogens in plant monocultures upon pesticide treatment or seasonal clonal turnover in *Daphnia* (McDonald et al., 2016; Steiner & Nowicki, 2019). However, this demographic scenario does not reflect the genetic starting conditions of more heterogeneous natural populations facing rapid environmental change. Adaptation dynamics from standing genetic variation differ substantially from the dynamics of isogenic populations. For instance, adaptation from standing genetic variation is more likely to lead to the fixation of more alleles of small effect relative to *de novo* mutations (Barrett & Schluter, 2008) imposing constraints on allele frequency dynamics through linkage, epistasis and pleiotropy (Hermisson & Pennings, 2005). Recently, evolution studies with yeast have allowed for more genetic diversity in the founder populations, and by inducing rounds of sexual reproduction over the course of experimental evolution (e.g., Burke et al., 2014; Kosheleva & Desai, 2018; Linder et al., 2020, 2022; Parts et al., 2011; reviewed in Phillips et al., 2020). However, environmental complexity is often extremely reduced in long-term microbial experimental evolution studies; experiments are either conducted in laboratory environments imposing no stress or focusing on the effects of a single stressor only.

Here, we address the impact of both of these aspects – genetic and environmental diversity – on adaptation dynamics, by using experimental microbial evolution with genetically diverse *S. cerevisiae* yeast populations in four environments with different stress levels. We tracked changes at the phenotypic and genetic level, driven by both standing genetic variation and *de novo* mutations. Founder populations were generated by crossing two strains of yeast (Y55 and SK1) that are 0.35% divergent genome-wide (Liti et al., 2009) and have high-quality genomic resources available. The F1 offspring was mass-sporulated to generate a diverse founder population of genetically unique, recombined F2 genotypes. Replicate populations of these founders thus contained millions of different genotypes initially (**Figure 1**) before we subjected them to selection in different environments for up to 1000 generations. The fitness of the evolved populations was compared to the fitness of the founder population at five time points of evolution (in ancestral and stressful environments), and populations were pool-sequenced at nine time points to track when and how the standing genetic variation of the founder was sorted. We explored the dynamics of ancestral and *de novo* allele frequencies, and also variation in chromosome copy numbers, because aneuploidies can increase the fitness of yeast in stressful environments (reviewed in Gilchrist & Stelkens, 2019). Our design allowed us to assess the degree of parallel molecular and phenotypic evolution in replicate populations selected in the same and different environments. We made four broad predictions: 1) Populations increase in fitness over time and fitness gains in stressful environments are larger, 2) the genetic diversity present in the founder will be rapidly sorted and decrease over time, 3) adaptation will be driven by standing genetic variation first and by *de novo* mutations later in the experiment, and 4) the degree of parallelism between replicates is larger in more stressful environments.

**Figure 1.**
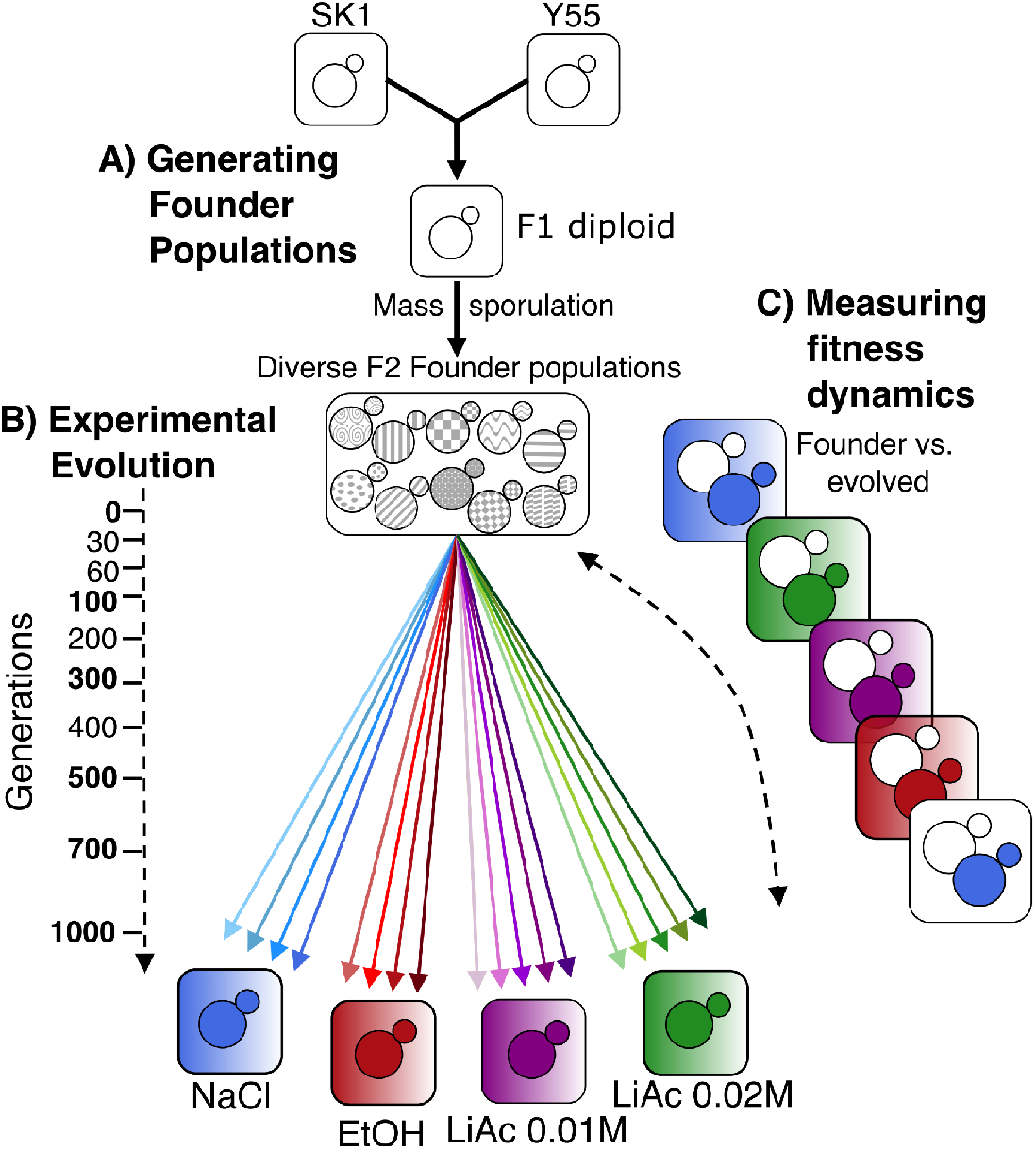
Experimental Design. **A)** Diverse founder populations were generated by mass sporulation of a cross between strains Y55 and SK1. **B)** Asexual yeast populations were evolved for 700-1000 generations in four environments: NaCl 0.75M, EtOH 8%, lithium acetate 0.01M (700 generations only), and lithium acetate 0.02M (700 generations only). Coloured squares indicate selective media; white squares are the ancestral medium (SC). Replicate populations are indicated as coloured arrows. Sampling timepoints for sequencing are indicated on the dotted arrow. **C)** Fitness assays using optical density (OD) after 24h of growth, comparing founders and evolved populations at 6 time points of evolution in selective and ancestral environments. Only a subset of all pairwise comparisons are shown. Assays were carried out at 6 time points of divergence (marked in bold on the left).

## Methods

### Strain engineering and crosses

We made F1 crosses between *S. cerevisiae* strains SK1 (*MATα ho::KANMx*) and Y55 (*MATa ho leu2::HYGMx*) and sporulated them in 50ml KAc media (1% potassium acetate, 0.05% glucose, 0.1% yeast extract made up to 1L with sterilised, deionised H_2_O) to generate large, genetically recombined diploid founder populations with standing genetic variation. To be able to identify populations on drop out media, we transformed the original SK1 and Y55 strains to contain auxotrophies for genes that synthesise essential amino acids. Following the protocol by Gietz and Schiestl (2007), we knocked out the functional copies of either uracil (ura3) or lysine (lys2) and replaced them with a drug resistance gene (nourseothricin sulphate; clonNAT). We verified the successful insertion of all drug cassettes by PCR. Founding strains thus contain alternative antibiotic resistance markers and auxotrophies, but otherwise the same strain background. Specifically, the founders of populations evolving in NaCl are the diploid F2 offspring from a cross between SK1 (*MATα ho:KANMx ura3::ClonNat*) and Y55 (*MATa ho leu2::HYGMx ura3::ClonNat*). All other populations (LiAc 0.01M, LiAc 0.02M and EtOH) are the diploid F2 offspring of a cross between SK1 (*MATα ho:KANMx lys2::ClonNat*) and Y55 (*MATa ho leu2::HYGMx lys2::ClonNat*). Note that NaCl adapted populations lost their ura3 auxotrophy within the first 100 generations, but maintained all other expected markers. The fitness of the founder populations was measured in selective environments (Optical Density or OD_600_ after 24h in culture) and used to standardise the evolved populations’ fitness.

### Environments and Serial Transfer

All environments consisted of synthetic complete (SC) medium (2% glucose, 0.67% bacto-yeast nitrogen base without amino acids, 0.00079% Formedium CSM powder (product code DCS0019) made up to 1L with sterilised, deionised water) plus a stress-inducing substance. We used four substances: lithium acetate (LiAc) 0.01M, LiAc 0.02M, sodium chloride (NaCl) 0.75M, and ethanol (EtOH) 8%. Replicate populations were grown in separate 60 ml glass test tubes containing 5ml of SC media with the appropriate stressor added. Every 48h, 10µl of culture were transferred to fresh media. This corresponded to around seven generations, estimated by counting cells after 48 hours growth in selective media with 0.75M NaCl. For the EtOH environment, the correct volume of EtOH was added to the tube just before inoculation to limit evaporation. Every five transfers (10 days, representative of ∼35 generations) 900µl population samples were frozen in 900µl of 44% glycerol and stored at -80°C for further analyses. Populations were checked regularly for environmental contamination by microscopic inspection and plating out on agar. We controlled for cross-line contamination using the selectable alternative genetic markers. The experiment was run for a total of ∼300 days.

### Fitness Dynamics

We measured the fitness as OD_600_ after 24h of growth in culture of all evolving populations at generations: 0, 100, 300, 500, 700, and 1000. All populations were measured in the environment they were selected in, and in the ancestral medium (SC only). Optical density was measured using a microplate reader (Sunrise, Tecan) using the accuracy mode at 600nm absorbance. To do this, frozen population samples were thawed and 10µl was used to inoculate 5ml SC media, then grown for 24h at 30°C with shaking. After 24h, 96-well plates were filled with 148µl per well of the appropriate medium and inoculated with 2µl of the 24h yeast culture. OD was measured at t_0_ and after 24h of growth at 30°C without shaking. Plates were gently vortexed for 30s and shaken in the plate reader for 20s immediately before OD readings. Each replicate population was measured in five wells and normalised using blank wells containing only media. Founder populations were measured separately, with 50 wells per environment in order to capture the potential phenotypic diversity present in the founding population. Relative yield for adapted populations was calculated as OD_adapted_/mean(OD_founder_) for each OD measurement taken. A value of >1 represents higher fitness than the mean founder population fitness. Raw fitness data is available in **Supplementary Table 1**.

Type II ANOVA and Tukey HSD tests were used to compare the fitness of evolved populations to the founder, as well as the founder’s fitness in the selective environments to fitness in unstressed conditions. To explain variance in fitness between replicate populations, we used a linear model with generation, replicate and their interaction as fixed effects, selecting the simplest model using type I ANOVA. All statistical analyses and graphs were done with R v4.1.0 (R core team, 2021), using the packages, car v3.0-5 (Fox & Weisberg, 2019), dplyr v1.0.7 (Wickham et al., 2021), emmeans v1.7.1-1 (Lenth, 2022), ggplot2 v3.3.5 (Wickham, 2016), gridExtra v2.3 (Auguie, 2017; Venables & Ripley, 2002) and MASS v7.3-54 (Venables and Ripley, 2002). Scripts used for analyses and plotting available at https://github.com/SLAment/AdaptationDynamics.

### DNA extraction and whole genome sequencing

DNA was extracted from whole population samples at the nine time points (0, 30, 60, 100, 200, 300, 400, 500, 700, and 1000 generations) of experimental evolution. For this, 10µl of frozen populations were grown in 5ml SC media at 30°C shaking at 200 rpm for 24-48h (until near saturation) and DNA was extracted using Thermo Scientific KingFisher™ Duo Prime Purification, as per manufacturer’s instructions (see supplementary materials for further details). Library preparation and sequencing were performed at the Science for Life Laboratories (Stockholm, Sweden). Library preparation followed the Illumina Nextera Flex protocol and sequencing was carried out by Illumina pool-seq using either the NovaSeq S4-300 (2x150bp) or NovaSeq S-Prime (2x150bp) to an estimated median depth of coverage per sample of around 30x to 780x. The raw sequences were deposited in the European Nucleotide Archive, accession number PRJEB46680. In addition, we included publicly available Illumina reads of the parental strains SK1 and Y55 in all analyses (accession numbers PRJNA340312 and PRJNA552112; Bendixsen et al., 2021; Yue et al., 2017).

### Quality control, mapping and variant calling

Both FastQC v0.11.9 (Andrews, 2010) and MultiQC v1.11 (Ewels et al., 2016) were used to assess the quality of the raw sequencing reads, both before and after trimming by trimmomatic v0.36 (Bolger et al., 2014). Samples LiAc0.02 R2 (replicate 2) at 60 generations and LiAc0.01 R5 at 500 generations were dropped due to poor coverage and sequencing failure respectively. The genome of strain S288C (R64; GenBank Accession GCA_000146045.2) was used as the reference. We then carried out alignment using BWA v0.7.17 (Li & Durbin, 2010) and samtools v1.9 (Li et al., 2009) to sort the generated bam files. Duplicated reads were removed using picard v2.23.4 (http://broadinstitute.github.io/picard/), followed by quality assessment of the mapping with QualiMap v2.1.1 (Okonechnikov et al., 2015) and MultiQC. To call variants, we used GATK v4.1.4.1 (McKenna et al., 2010) HaplotypeCaller best practices, by first generating g.vcf files, followed by joint calling of all samples. SNPs and INDELs were extracted and filtered separately using the following cut-offs: SNPs – QD<2.0, FS > 10.0, ReadPosRankSum <-3.0, MQRankSum < -6.0, SOR > 3.0, MQ < 40.0; INDELs – QD <10.0, FS > 10.0, ReadPosRankSum <-4.0, MQRankSum < -8.0, SOR > 4.0, MQ < 40.0. Subsequently, we used two filtering strategies to recover high quality biallelic variants, one to characterize standing genetic variation across all environments, and another to identify *de novo* mutations exclusive to each environment. For the first strategy, we removed sites with missing data using the option *--max-missing* 1 in vcftools v0.1.16 (Danecek et al., 2011), and further filtered out sites where at least one sample had coverage below 25x or above the 95th coverage sample percentile in R, as well as sites that had a mappability of less than 1 as calculated by GenMap 1.3.0 (Pockrandt et al., 2020). From these filtered variant set, we selected sites that showed evidence of being homozygous (minor allele frequency, MAF < 0.05) in the haploid parental strains SK1 and Y55 (false heterozygous sites would be produced by mismapping and hidden paralogy) and that were also clearly heterozygous in the Founder samples (MAF > 0.3), which by design should be exclusively F2 heterozygotes. From these, the final variant set contained only sites that were shared across all samples in all environments. For the second strategy, we first split the variants by environment using vcftools and remove sites that became monomorphic (based on allele depth) using the custom script *getvariantspool.py* available in the GitHub repository (see below). We then removed sites with missing data, bad coverage and mappability, and heterozygosity in the parental strains as above. Finally, we used snpEff v5.0e (Cingolani et al., 2012) to annotate the effect of identified variants on genes using the database of the reference genome R64-1-1.99 with canonical transcripts (*-canon*). The workflow is available at https://github.com/SLAment/AdaptationDynamics as Snakemake v. 5.30.1 (Köster & Rahmann, 2012) pipelines, which depend on the ggplot2 v. 3.3.3 (Wickham, 2016), cowplot v. 1.1.1 (Wilke, 2019), dplyr v. 1.0.5 (Wickham et al., 2021), vcfR v. 1.12.0 (Knaus & Grünwald, 2017), and ggpubr v. 0.4.0 (Kassambara, 2020) R packages.

Preliminary analyses revealed indications of contamination in a few samples. Namely, in some samples the majority of reads mapped to another yeast species also present in the lab environment (*Saccharomyces paradoxus*) or multiple samples shared sets of *de novo* mutations (detected as described below) that appeared simultaneously in the last time points of some populations, while at the same time exhibiting a sudden increase in allelic diversity across the genome. As we deemed these observations unlikely to evolve naturally in multiple populations simultaneously, we excluded two NaCl-adapted populations after 700 generations.

### Analysis of standing genetic variation

Due to our experimental design, in which only one round of sexual recombination occurred, we expect that large haplotypes were formed in the initial recombination event. To assess the dynamics of these haplotype blocks, we implemented a method of reconstructing selected haplotypes based on correlation between SNPs through time, inspired by Otte & Schlötterer’s Haplovalidate (Otte & Schlötterer, 2021). From the set of SNPs filtered with the first strategy as described above, we first identified subsets of SNPs likely to have been affected by direct or, more likely, indirect selection. We estimate the variance effective population size between every pair of available timepoints for each replicate with the *estimateNe* function of the poolSeq (v. 0.3.0) R package (Taus et al., 2017). Within the function, *Ncensus* and *poolSize* were set to 1,000,000 and 100,000, respectively. These metrics are likely underestimating the actual *N* and pool size, though increasing them further does not change estimated Ne results. Given the estimates of Ne, we then ask whether the change in allele frequency for each SNP deviates from expectations of a null drift model. We calculated the probability of the observed allele frequency change again between every pair of timepoints based on the beta approximation to the Wright-Fisher model (Ewens, 2012; Gompert & Messina, 2016), and retain SNPs with allele frequency changes more extreme than the 1st or 99th percentile of the null distribution for any of the intervals. To reconstruct haplotype blocks, we first normalized allele frequency data of the reduced set of SNPs with an arcsine-square-root transformation with subsequent centering and scaling (Otte & Schlötterer, 2021). We then calculated correlations between all SNPs and performed hierarchical clustering to obtain haplotype blocks. Only haplotype blocks with at least 10 SNPs were retained. We tested various minimum correlation cutoffs for hierarchical clustering from 0.3 to 0.9, and chose the correlation cutoff for each replicate which resulted in the most haplotype blocks of minimum size. We chose one SNP at random within each haplotype cluster to represent the overall cluster trajectory. Notice that each haplotype cluster might include multiple chromosomes, since reproduction in our experiment is strictly asexual. This aspect of our data also makes the original HaploValidate algorithm, which relies on linkage equilibrium between chromosomes, non-applicable to our experiment.

The original parental strains, SK1 and Y55, have a nucleotide identity percentage of 0.35% (Liti et al., 2009), but preliminary analyses revealed that the divergence between these strains is not uniform along the genome. Instead, the divergence between them is distributed in blocks, with some large chromosomal regions being identical between strains. Thus, our analyses of standing genetic variation below include relatively few sites of some chromosomes (e.g., chromosomes 2, 6 and 10).

### Analysis of de novo mutations

To identify *de novo* mutations, we compared the alleles of the parental strains SK1 and Y55 with alleles found in the biallelic sites of the evolved samples in the second variant dataset. All sites that had an allele not found in any of the parental strains and that had a MAF of less than 0.3 in the founder populations were considered putative *de novo* mutations. The MAF requirement excludes a few mutations (29 SNPs and four INDELs) that presumably appeared between the sequencing of the parental strains and the sequencing of the founder populations, which were already at intermediate frequencies (equal or more than 0.3) at the beginning of the experiment. Some of these intermediate mutations could be the result of read mismapping and/or paralogs and faulty variant calling. As many of the remaining putative mutations are also either sequencing errors or the result of mismapping as above, we further filtered for mutations that reached a frequency of 10% at least once in the experiment and considered them *de novo* mutations (310 variants). From this, we identified candidate adaptive mutations by filtering for variants that reached a frequency larger than 35% in at least one sample. The resulting 92 variants were subject to manual curation by examining mapped reads in the Integrative Genome Viewer v. 2.12.2 (Robinson et al., 2017). Sites with reads mapping to multiple locations, variant caller inconsistencies, and sharp increases in coverage were discarded (30 variants). Notice that the large proportion of discarded *de novo* mutations does not reflect the overall quality of the total variant set. Instead, it reflects the fact that bad quality sites have a tendency to create high-frequency variants. Out of the final 62 curated mutations, 57 were located within protein-coding genes. We used the list of these genes as input for the STRING database (Szklarczyk et al., 2021) to infer functional associations, last consulted on March 7th 2022 with default parameters.

We calculated the probability of mutations affecting the same gene independently as follows (Huang et al. 2018):

*(Freq. of mutations per gene in lineage 1)*(Freq. of mutations per gene in lineage 2)*Total No. of genes* We assumed the total number of genes to be 6427 as in the S288C reference genome. As an example, consider the gene *CYC8*, which had two independent mutations in replicates 2 and 3 of the LiAc 0.02M environment. As there were five and four high frequency (> 0.35), manually curated mutations in these two replicates at the end of the experiment, respectively, the probability of observing two independent mutations in a gene of these two replicates is: (5/6427) x (4/6427) x (6427) = 0.0031. Notice however that a total of 6427 genes leads to an underestimation of this number because many sites across the genome will not be callable due to issues with coverage and mismapping. Still, even using as few as 1000 genes leads to probabilities lower than 0.03 for all genes with multiple hits (not shown).

### Analysis of copy number variants and potential aneuploidies

To identify potential aneuploidies, we ran samtools depth only counting reads with mapping quality greater than or equal to 20 (-Q 20) for each basepair across the genome. Then each replicate population was analysed using a custom python pipeline, which determined the mean read depth in 1kb non-overlapping windows, removed outliers (>2x mean chromosome read depth), determined mean read depth within each chromosome and calculated the mean read depth across the genome for the replicate population. The depth of each chromosome was then compared to the genomic mean depth, and was characterised as a potential aneuploidy in the population if chromosome depth was +/-25% of the mean genome depth. For a few sequenced time points, read depth was not uniformly distributed along chromosomes, with chromosome centres having less read depth, resulting in a noticeable ‘smile’ shape in read depth. The cause of the unusual distributions is unknown, but was not linked to overall sample read depth or to sequencing flow-cell. For time points where read depth was highly variable within chromosomes (standard deviation greater than 15% of mean read depth and chromosome centre more than 15% lower than chromosome ends), the average chromosome-end read depth (excluding sub-telomeric regions) was used as the chromosome mean. These were calculated individually for each replicate/generation time point, and plotted using custom python scripts.

### PCR verification of mating type

Populations can change ploidy and mating type over time. While all populations in this experiment were *HO* knock-outs (they cannot switch mating type and self-fertilise), ploidy changes may still occur. In order to test the ploidy of our populations, we used PCR primers designed to target within the mating loci genes (Illuxley et al., 1990) in generations 30, 60, 100 and 200 from the LiAc 0.02M environment. Primers were designed to display a band at 492bp for the MATa or 369bp for the MATα allele. Two bands on the gel indicate a non-haploid population (likely diploid as in the founder populations).

## Results

### Fitness dynamics

We predicted that selective environments with higher fitness cost have stronger impacts on fitness dynamics and allele frequency changes. To test this, we assessed the fitness cost of each environment by comparing the fitness of the founder populations in each selective environment to their fitness in ancestral conditions (SC media). Selective environments significantly differed in how much they reduced the fitness of the founder populations (**Figure 2A**; NaCl founder: F_118_ = 3826.2, *p* < 0.001; LiAc and EtOH founder: F_236_ = 286.7, *p* < 0.001). NaCl 0.75M (t = 61.9, df = 118, *p* < 0.001) and LiAc 0.02M (t = 25.7, df = 236, *p* < 0.001) had the highest fitness costs. LiAc 0.01M showed a more modest cost (t = 5.5, df = 236, *p* < 0.001). Ethanol 8% had no significant impact on fitness (t = 1.1, df = 236, *p* > 0.05).

**Figure 2.**
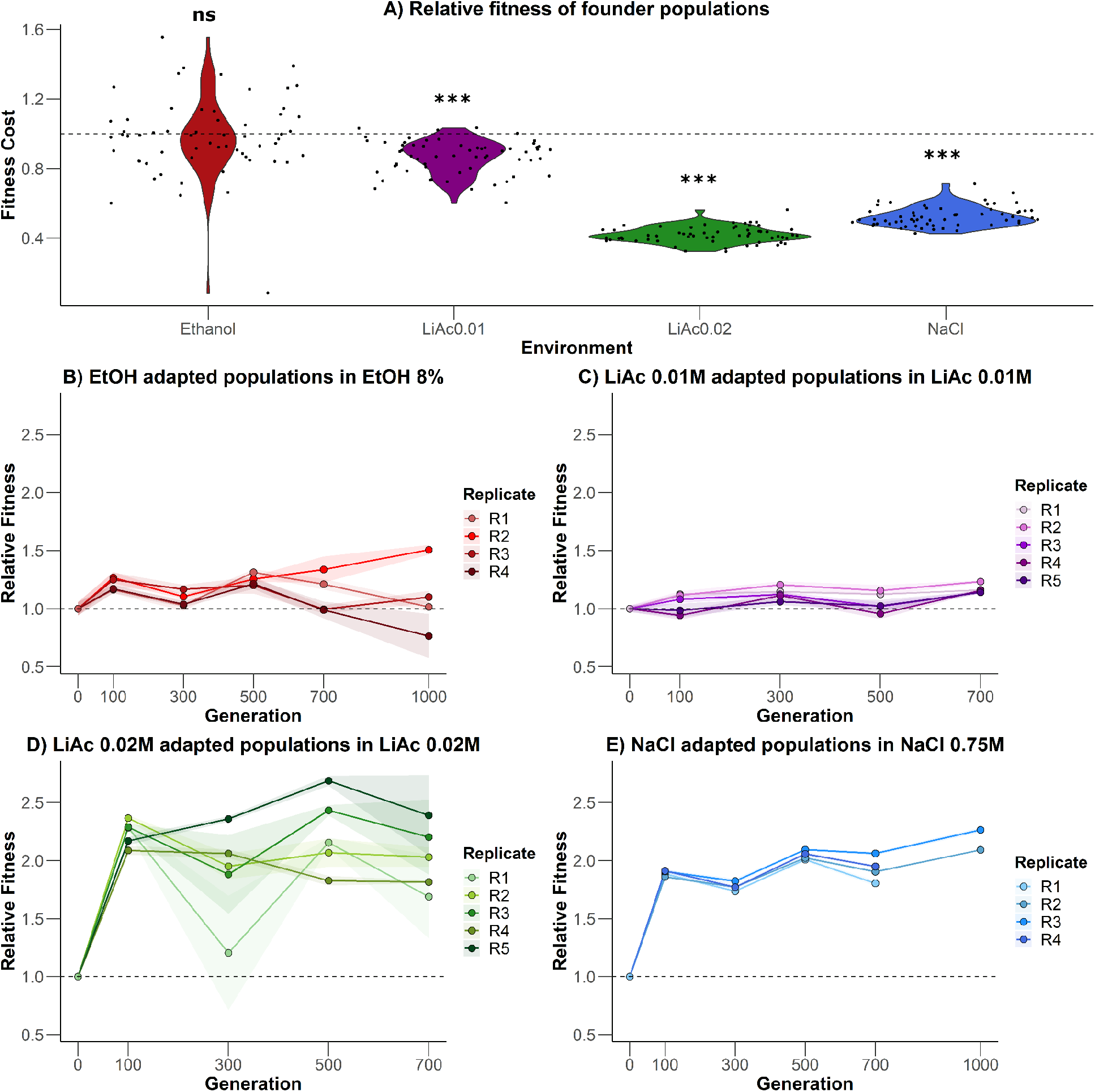
Fitness dynamics. **A)** Founder population fitness (OD600 after 24h of growth) in selective environments compared to growth in ancestral conditions (SC medium). Values of 1 (dashed horizontal line) indicate no change compared to unstressed conditions. Values below 1 indicate lower fitness in selective media, i.e. a fitness cost. Each wee black point represents a single measurement. **B-E)** Mean relative fitness of replicate populations in the four selective environments. Shaded areas represent the 95% confidence intervals for each replicate population. One EtOH population went extinct in the first 100 generations. Two NaCl populations after 700 generations were removed due to contamination. *** = p < 0.001; ns = not significant.

Next, we tested if evolving populations increased in fitness in their selective environments over time. Variation in fitness between replicate populations within environments was mainly explained by an interaction between replicate and generation, i.e. fitness showed different trajectories over time in different replicates (**Figures 2B-E**, all F > 3.48, *p* < 0.001, df = 21, 308 for NaCl, df = 23, 314 for EtOH, and df = 24, 375 for both LiAcs). By the end of the experiment (700 generations for LiAc 0.01M, LiAc 0.02M and NaCl R1 and R4, and 1000 generations for EtOH and NaCl R2 and R3), all replicate populations in the NaCl, LiAc 0.01M and LiAc 0.02M environments had significantly higher fitness than the founder (all pairwise Tukey HSD tests t > 3.48, *p* < 0.01, df = 308 for NaCl, df = 375 for both LiAcs). On average, fitness increased by 118% in NaCl, 17% in LiAc 0.01M, and 103% in LiAc 0.02M. Replicate populations adapting to the two most stressful environments (NaCl and LiAc 0.02M) had significantly higher fitness than the founder at all generations tested (t > 2.78, *p* < 0.05). This was not the case for populations evolving in the lower stress environments (EtOH and LiAc 0.01M). Three of the four replicate populations evolving in EtOH did not significantly differ from the founder at the end of the experiment (R1, R3 and R4: t < 2.43, df = 314, *p* > 0.1; R2: t < 6.42, df = 314, *p* < 0.001). In general, significant differences between replicate populations within each environment increased at later generations (mean of 15.8% significant pairwise Tukey HSD tests at 100 gens vs. 48.3% significant pairwise at 700 gens for all possible comparisons).

Averaged across replicates, the largest single increase in fitness occurred early on in three of the four environments (**Figures 2B-E**), that is between 0 and 100 generations (75.9% of total in NaCl, 89.3% of total in EtOH, and 100% in LiAc 0.02M). In LiAc 0.01M, fitness only increased by 29.7% in the first 100 generations. When tested in ancestral conditions (SC medium), evolved populations showed significantly higher fitness than the founder populations by the end of the experiment, although the magnitude of increase was small for LiAc environments (F_21, 718_ = 251.77, *p* < 0.001; **Supplementary Figure 1**). In ancestral conditions, NaCl populations showed the largest increase in relative yield within the first 100 generations, while populations in EtOH increased only at later generations. This increased fitness in ancestral conditions could indicate adaptation to the SC component of the stressful environment (Johnson et al., 2021) or suggest that alleles providing high fitness in stress environments are also beneficial in ancestral conditions.

### Allele frequency dynamics

In order to compare between replicate populations and environments, we selected a set of 9940 SNPs originally present in the two parental strains and shared across all samples (i.e., no missing data). As expected from genetic drift and selection, we observed a tendency of alleles to either go to fixation or extinction as time progressed in the experiment (**Supplementary Figures 2-5**). Typically, a large proportion of ancestral variants settled into intermediate frequencies (around 0.5) suggesting the fixation of a single diploid genotype. Sets of variants shared similar allele frequency trajectories (Supplementary **Figures 2-5**). Hence, we reduced the 9940 SNPs to sets of correlated variants or haplotype clusters in order to represent the overall ancestral variation dynamics (**Figure 3**). Since we evolved F2 genotypes that underwent only one round of recombination, genetic linkage is high across the entire genome and remains high during the whole experiment. Naturally, there are cases where a haplotype contains a mix of the parental Y55 and SK1 alleles. Because they were not recombined off that background, this results in allele frequency trajectories mirroring each other (**Figure 3; Supplementary Figures 2-5**), due to an increasing Y55 allele appearing as a decreasing SK1 allele, and vice-versa.

**Figure 3.**
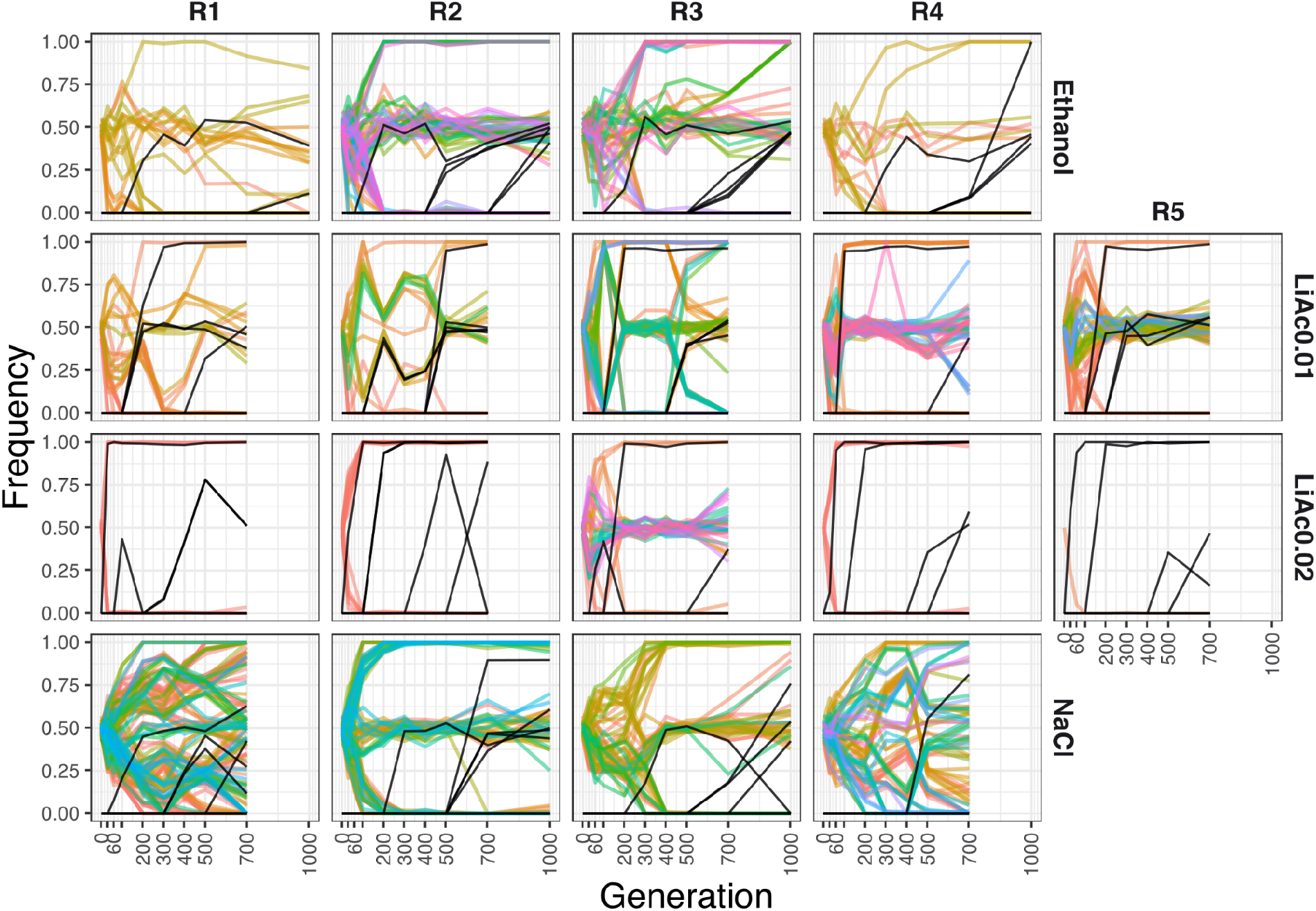
Frequency trajectories. Haplotype clusters of standing genetic variation (in colour) and high-frequency de novo mutations (black) across environments (rows) and replicates (columns). The allele frequency of the haplotype clusters is based on the SK1 allele.

**Figure 5.**
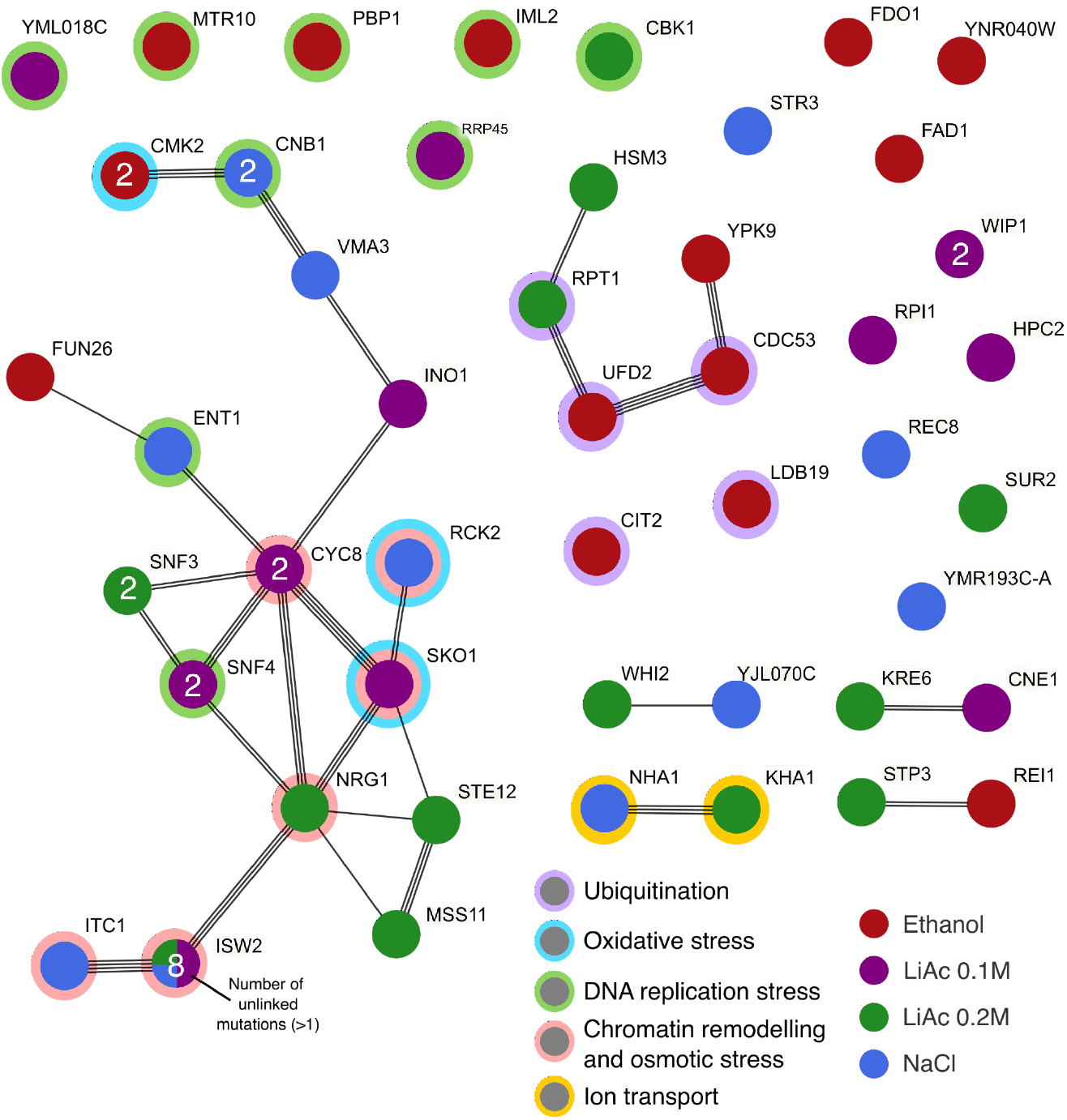
Protein–protein interaction network of all genes affected by *de novo* mutations that reached frequency of at least 35% at some point during the experiment. Each node is a pie chart of the proportion of mutations per gene that occurred in each environment (all present in a single environment, except for *ISW2*). The number of unlinked independent mutations is indicated within each node if a gene was hit by more than one mutation (two mutations in the genes *SUR2* and *SNF3* have linked allele frequency trajectories and were counted as one). The links represent different sources of interaction evidence detected by the STRING database. Main functional categories are depicted by rings around the nodes.

While each environment had its own idiosyncrasies, the LiAc 0.02M environment showed the most drastic changes in allele frequencies (**Figure 3**; **Supplementary Figures 2-5**). Specifically, in four out of five LiAc 0.02M replicates, we observed that the standing genetic variation was depleted extremely quickly, leaving no intermediate allele frequencies. This pattern is consistent with the fixation of a single dominant genotype that was haploid or a fully homozygous diploid. In line with this interpretation, PCR amplification recovered a single mating type (*MATα*) already from generation 30, the earliest time point tested, in all replicates apart from replicate 3.

Regardless of the drastic changes of some haplotypes, a considerable fraction of variants reached fixation in all populations. Fitting a sigmoidal model to the proportion of nearly fixed sites in each replicate population shows that the curves often plateau at proportions close to 0.5, 0.75 or 1 (**Figure 4A**). A proportion of 0.5 is expected if a single F2 genotype was fixed in a given population, since their inbreeding coefficient is ½ by experimental design (equivalent to one round of intertetrad mating; Kirby, 1984). The proportion of 1 is consistent with a single haploid or fully homozygous diploid going to fixation. The origin of the 0.75 fraction is less clear. Overall, the maximum proportion of nearly fixed sites differs between environments (Kruskal Wallis H_3_ = 11.18, *p* = 0.011), mostly due to the complete fixation of alleles in the LiAc 0.02M environment (**Figure 4B**). Plateauing also occurred faster in LiAc 0.02M, typically within the first 100 generations, than in any other environment (**Figure 4C**; Kruskal Wallis H_3_ = 12.09, *p* = 0.007).

**Figure 4.**
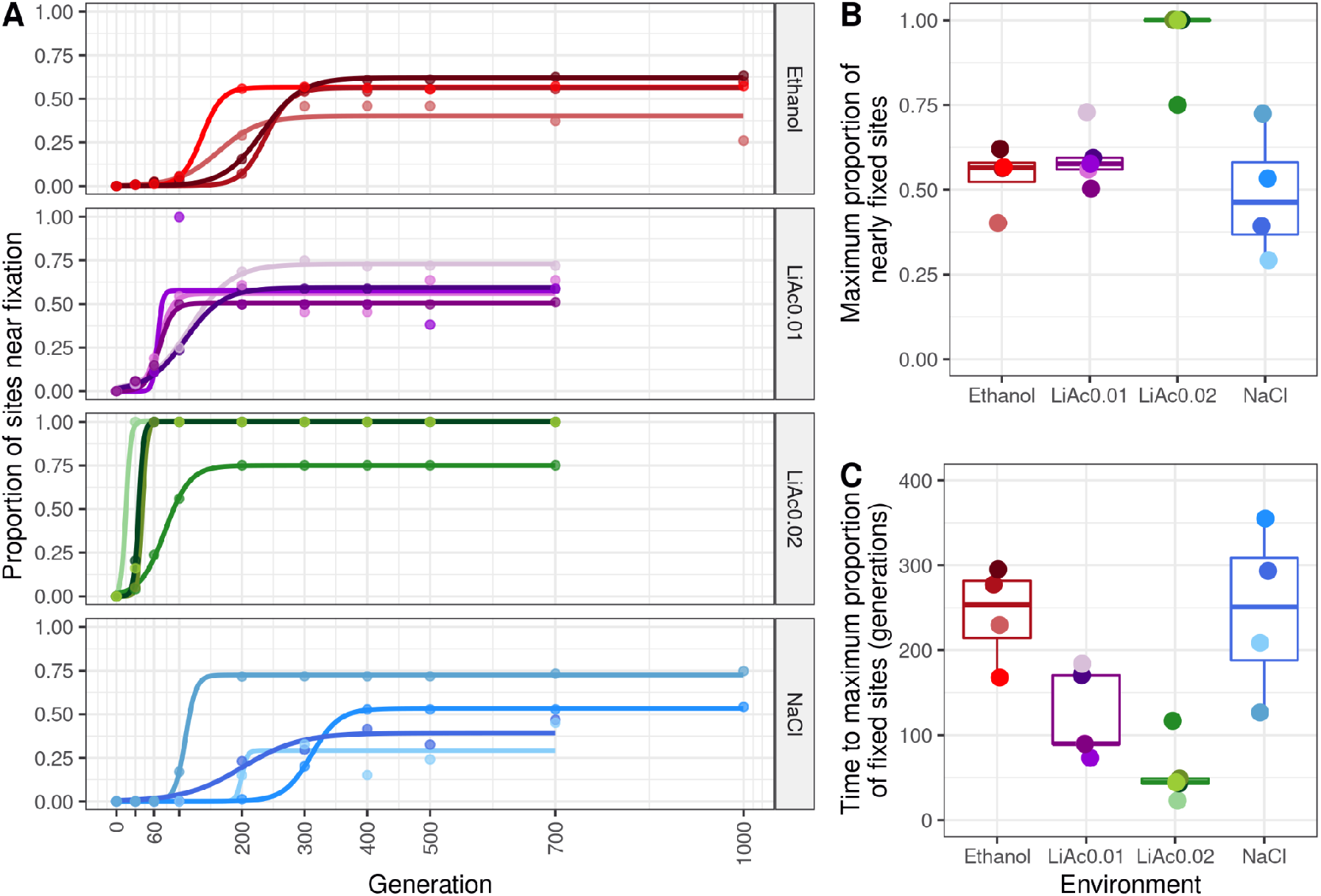
Differences in the proportion of nearly fixed sites (MAF < 0.1) across environments. **A)** The raw proportion of nearly fixed sites through time (points) were fitted to a sigmoidal model for each replicate, **B)** each with their respective maximum proportion of nearly fixed sites, and **C)** the time at which each replicate reaches this maximum.

These observations are consistent with stronger selection in LiAc0.02. Although the NaCl environment also imposed high fitness costs (**Figure 2A**), there was large variation between replicates in the proportion of nearly fixed sites, and diversity was generally lost at a slower pace than in the other environments (**Figure 4C**; but pairwise Wilcoxon tests were not significant after Bonferroni correction). Likewise, many distinct haplotypes were recovered in replicates 1 and 4 of the NaCl environment (**Figure 3**), suggesting that multiple genotypes persisted for longer in those populations.

Surprisingly, the direction of change in the allele frequency of some variants shifted during the duration of the experiment (**Figure 3**; **Supplementary Figures 2-5**). In extreme cases, some alleles approached fixation at a given time point but then returned to intermediate frequencies (e.g., in the replicate 3 of LiAc 0.01M; **Figure 3**). To further explore this pattern, we calculated the proportion of sites near fixation (MAF < 0.1) for each sample (every time point of each population). We found that the proportion of nearly fixed sites at a given time point is not explained by the depth of coverage (**Supplementary Figure 6**), suggesting that the “unfixing” of sites is not an artefact of sequencing. However, some drastic frequency changes did coincide with the trajectory of *de novo* mutations (**Figure 3**), which in principle could change the direction of selection for the haplotype containing them (see next section for further discussion).

### *De novo* mutations

From the detected 310 putative *de novo* mutations (183 SNPs and 127 INDELs), we focused on potentially adaptive variants that reached a frequency larger than 35% during at least one time point in any sample of the experiment (56 SNPs and 6 INDELs after manual curation) (black lines in **Figure 3**). The SnpEff program predicted moderate to high fitness effects for these high-frequency variants, as expected if they were under selection (**Supplementary Figure 7**). Most of these *novo* mutations were located within protein coding regions (57 out 64), affecting in total 47 genes with six synonymous mutations, 35 nonsynonymous point mutations, six frameshift mutations, nine stop codon gains, and one start codon loss. Notably, a STRING network of these 47 genes has significantly more interactions than expected given a random set of proteins of the same size and node degree distribution (protein-protein interaction enrichment *p* = 0.00156; **Figure 5**). In particular, multiple independent mutations hit members and known cofactors of the protein complexes Isw2p-Itc1p and Cyc8(Ssn6)p-Tup1p, in almost all replicates of the NaCl and LiAc environments (red rings in **Figures 5**), reaching frequencies of around 0.5 (expected if a single heterozygous genotype fixed) or 1 (**Figures 6**). The transcriptional corepressor complex Cyc8p-Tup1p recruits the chromatin remodelling complex Isw2p-Itc1p to nucleosomes, controlling the repression of hundreds of genes under unstressed conditions (Green & Johnson, 2004; Rizzo et al., 2011; Zhang & Reese, 2004). Many of these genes are activated in response to cellular stress, including osmotic stress (Green & Johnson, 2004; Proft & Serrano, 1999; Proft & Struhl, 2002).

**Figure 6.**
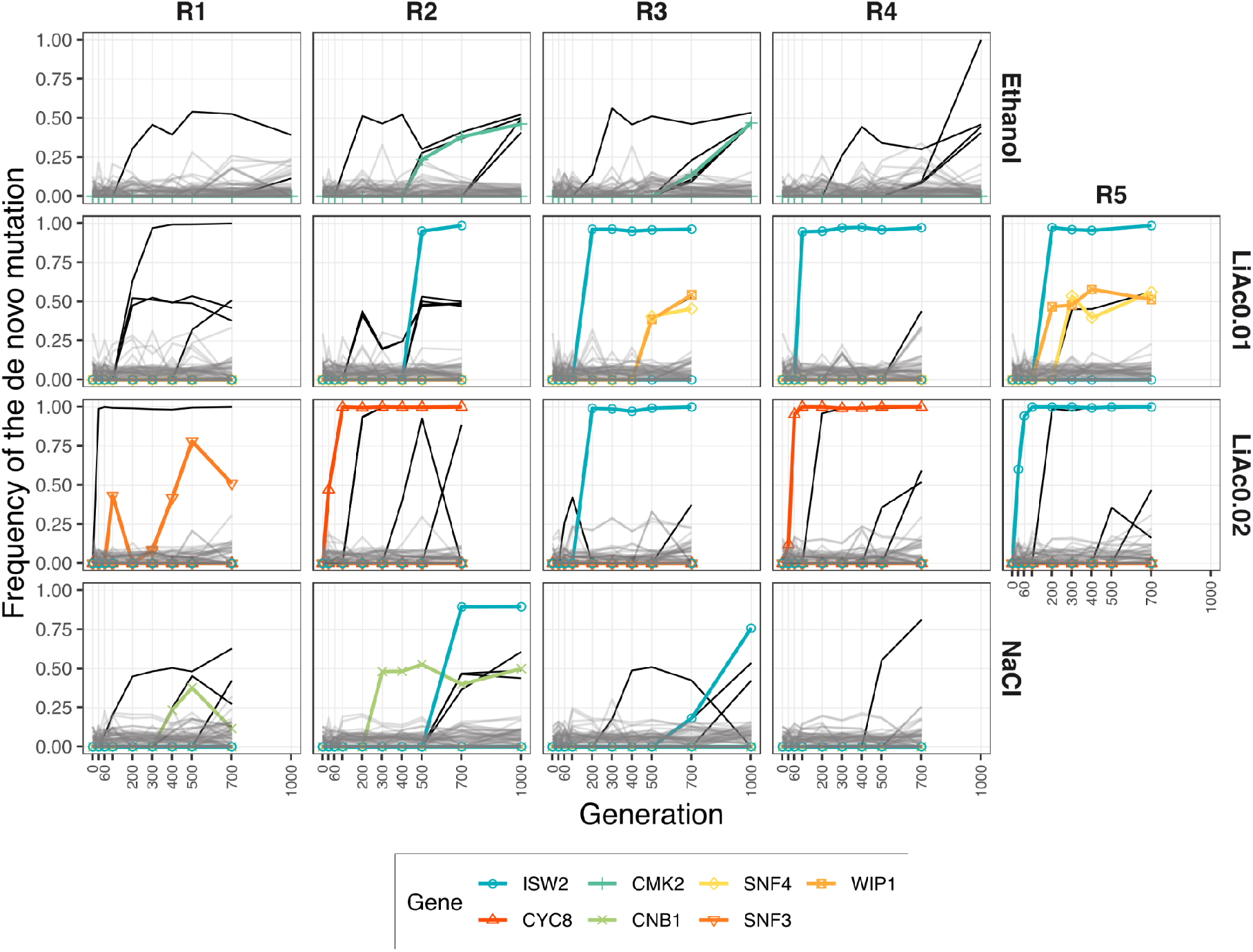
Genes with parallel independent *de novo* mutations. Of the *de novo* mutations that reached a frequency higher than 35% at some time point (black lines), some hit the same gene independently (in colours). Mutations that stayed below 35% frequency are presented in grey.

In our experiment, the gene *CYC8* (YBR112C or *SSN6*) gained two independent missense mutations in different replicas of LiAc 0.02M (**Figure 5**). Notably, it has been shown that the Cyc8p*-*Tup1p complex acts as a repressor of *SKO1* (YNL167C), a gene that gained an internal stop codon in one replicate of LiAc 0.01M (**Figure 5**). *SKO1*, in turn, is a downstream effector of the High-Osmolarity Glycerol (HOG) pathway, which activates during osmotic stress and glucose starvation (Proft & Serrano, 1999). Along with the Cyc8p-Tup1p complex, Sko1p represses the primary sodium and lithium-ion transporter *ENA1* (Proft & Serrano, 1999). Likely, deactivation of these members of the HOG pathway makes the expression of *ENA1* constitutive (at least to a degree, see Proft & Serrano, 1999), allowing for the constant expression of the genes involved in tolerating osmotic stress. Moreover, Sko1p is involved in recruiting other salt stress defence genes. Accordingly, deletions of *SKO1* have been shown to provide resistance against Na^+^ or Li^+^ (from NaCl and LiCl) stress (Proft & Serrano, 1999).

It is unknown if Isw2p-Itc1p interacts with Cyc8p-Tup1p while specifically repressing *SKO1*, but this is a strong possibility given that their effects on nucleosome positioning is highly correlated across the genome (Rizzo et al., 2011). Remarkably, up to eight independent *de novo* mutations disrupted the gene *ISW2* (YOR304W) in multiple environments (**Figure 6**), including the loss of the start codon, gains of stop codons and frame shifts. Likewise, one additional frame-shift INDEL affected the gene *ITC1* (YGL133W) in a NaCl population. Together, Isw2p-Itc1p are also known to repress early meiotic genes during mitotic growth (Goldmark et al., 2000). The de-repression of these early meiotic genes by deleting *ISW2* does not cause slow growth in cells (Goldmark et al., 2000). Hence, disruption of this protein complex might be advantageous during asexual reproduction under osmotic stress.

Similarly, in the NaCl environment, two nonsynonymous mutations affected the gene *CNB1* (**Figures 5** and **6**) that codes for the regulatory subunit of calcineurin, a regulator of *ENA1* transcription in response to sodium and lithium toxicity (Mendizabal et al., n.d.; Ruiz et al., 2003). Thus, the regulation of *ENA1* seems to be a common denominator in the molecular basis of adaptation to osmotic stress in our experiment. Of note, *ENA1* is part of a tandem repeat that leads to poor read mapping, which prevents detection of any mutations that could potentially affect this gene or its paralogs directly.

In addition, other genes affected by mutations in the NaCl and LiAc environments have functions associated with oxidative stress, ion transport, and DNA replication stress (**Figure 5**). By contrast, the high-frequency mutations found in the EtOH environment had less connections to those found in other environments, and were mostly associated with ubiquitination and DNA replication stress functions (**Figure 5**), which is in line with previous molecular characterization of the ethanol stress response (Navarro-Tapia et al., 2018; Stanley et al., 2010).

The trajectory of some mutations on their way to fixation changed abruptly. In fact, some mutations went extinct, precisely when another mutation appeared in the population (e.g. replicate 2 of LiAc 0.02M or replicate 3 of NaCl in **Figure 6**), in a pattern reminiscent of clonal interference (Gerrish & Lenski, 1998). This putative turnover of genotypes happened after the standing genetic variation was completely sorted in some populations, but it might have also affected the fate of standing genetic variation (e.g. replicate 2 of LiAc 0.01M in **Figure 5**). While it is tempting to speculate that some radical changes in the direction of a particular haplotype were due to the evolution of adaptive mutations, some mutations may in fact be neutral or deleterious and increased in frequency through genetic hitchhiking on a haplotype affected by other processes (e.g. cryptic fluctuating selection during the experiment). However, the fact that multiple high-frequency independent mutations hit the same genes in different replicates argues for their adaptive value (Kryazhimskiy et al., 2014). Specifically, the genes *CMK2*, *CNB1*, *CYC8*, *SNF3*, *SNF4*, and *WIP1* all had two unlinked mutations in two different replicates of different treatments, while the gene *ISW2* had eight (**Figure 6**), as mentioned above. The probability of multiple independent mutations affecting these genes is very low (*p* < 0.005, see Methods), and vanishingly small for *ISW2* in particular (*p* = 8.48 x 10^-23^; **Supplementary Table 2**).

### Chromosomal copy number changes

Using read depth, we found that changes in chromosomal copy number within the evolving populations were common over the course of experimental evolution. Thirty-nine percent (seven of 18) of all replicate lines contained at least one time point in which at least one chromosome deviated from the genome-wide mean read depth by more than 25% (**Supplementary Figure 8, Supplementary Table 3**). This was dominated by chromosome gains (89%) as opposed to losses (11%). While some of these potential aneuploidies were limited to single time points, we found several cases where aneuploidies were maintained in the evolving populations across hundreds of generations (**Figure 7**). For instance, in EtOH we detected the same aneuploidy (2 x chromosome 10 and 1 x chromosome 11) in at least three subsequent timepoints spanning 500 generations, which were maintained in the populations until the end of the experiment. Two notable examples of parallel patterns in copy number variation are chromosome 10 in EtOH and chromosome 3 in LiAc 0.01M, each of which significantly increased in relative chromosome read depth in two independent replicates (**Supplementary Figure 8**). No significant deviations were found in the NaCl environment. In addition, none of the *de novo* mutations that went to full fixation (frequency of 1) coincide with detected chromosome losses, suggesting that 1) the selective advantage of the detected aneuploidies might depend on the standing genetic variation, and 2) other mechanisms like gene conversion and meiotic recombination were involved in the fixation of *de novo* mutations that appeared in a diploid background.

**Figure 7.**
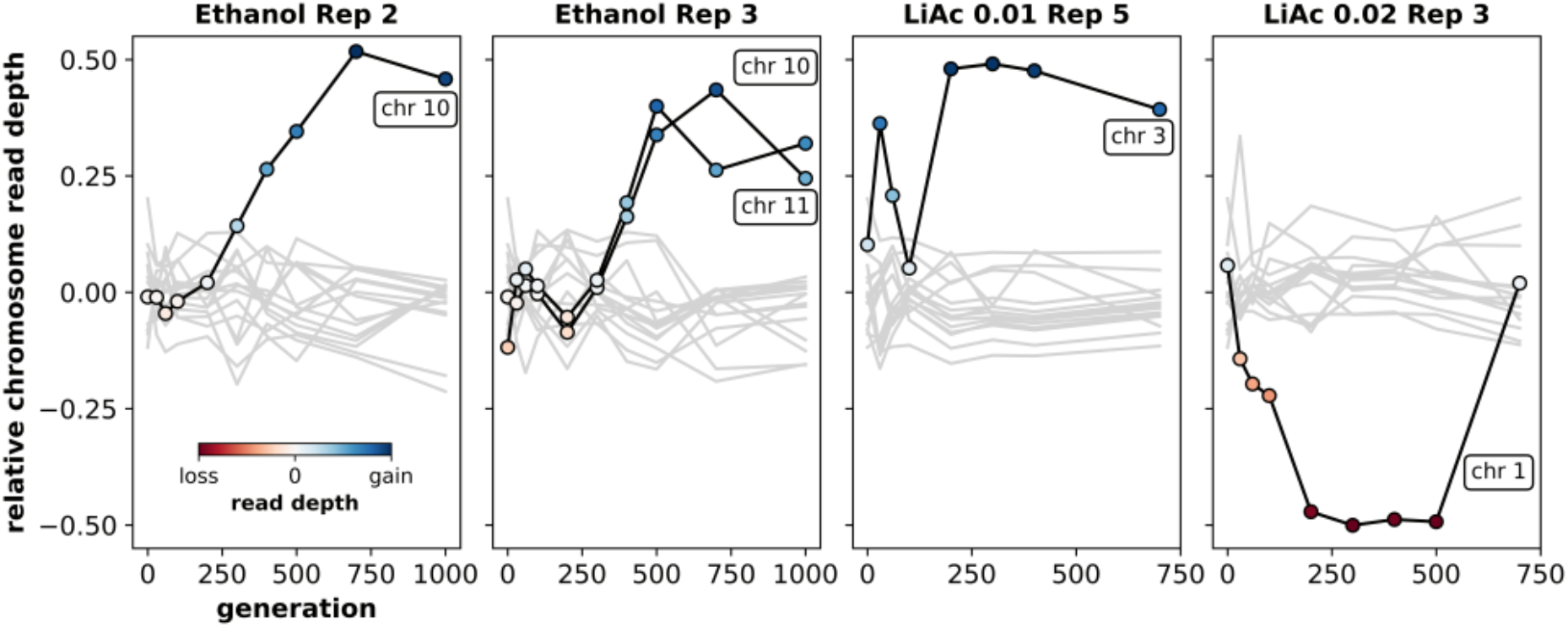
Chromosome copy number changes. A subset of chromosome copy number changes in four replicate populations. Lines show the relative read depth of each chromosome (read depth vs. the mean read depth across all chromosomes). Lines in black indicate chromosomes with potential aneuploidies. The points within indicate a gain (blue) or loss (red) of chromosome, with the shade indicating the magnitude of relative read depth change.

## Discussion

While adaptation from standing genetic variation is an important process underlying evolution in natural populations, we are often unable to characterise the timing and dynamics of fitness changes, and the underlying genetic mechanisms. Here, we used the power of microbial experimental evolution and whole population sequencing to track the phenotypic and genomic changes of genetically diverse yeast populations in environments with different stress levels. To simulate adaptation dynamics in more natural demographic conditions, we used outcrossed founder populations containing more standing genetic variation than the isogenic starting populations traditionally used in microbial experiment evolution. As expected, populations increased in fitness over time and showed significantly larger fitness gains in more stressful environments. The genetic diversity present in the founder was rapidly sorted and decreased quickly in most but not in all environments over time. Some populations of one environment in particular (NaCl) maintained multiple genotypes until the end of experimental evolution without the majority of alleles getting fixed. We detected parallelism at both phenotypic and genotypic levels (involving genes, pathways and aneuploidies) within and between environments, with conspicuous changes recurring in the high-stress environment LiAc 0.02M. In summary, our results suggest that adaptation was driven by both standing genetic variation and *de novo* mutations in our experiment, with interesting environmental idiosyncrasies.

### Adaptation dynamics in stressful environments

Populations normally increase in fitness over time in experimental evolution studies (e.g. Lenski et al. 1991, Johnson et al. 2021). We observed the strongest increases in fitness (> 100%) in the two environments with the high fitness cost (NaCl and LiAc in **Figure 2**) but found no or low fitness increases in less stressful environments. Most of the fitness increase occurred early in the experiment (in the first 100 generations), likely due to the sorting of the genetic diversity present in the founder. These fitness patterns are consistent with declining adaptability and diminishing returns of beneficial variants that have consistently smaller effects in fitter backgrounds (Johnson et al., 2019; Kryazhimskiy et al., 2014; Wiser et al., 2013). Environments with higher fitness costs likely contained genotypes with lower starting fitness, leaving more possibility for fitness improvements.

By the end of the experiment, many populations were dominated by a single diploid genotype, typical of populations evolving under strict asexual reproduction. The number of nearly fixed sites and the speed at which these sites reached near fixation increased with the stress level of the environments (**Figure 4**). One exception here is the NaCl environment, where we found more variation between replicates, alleles took a longer time to fixation, and the number of alleles nearing fixation was lower than in the other high stress environment. It is possible that this reflects the complexity of the trait, suggesting that adaptation to salt is polygenic and involves many loci of small effect with interactions between them. On the other hand, *de novo* mutations occurred in similar pathways in both NaCl and LiAc environments, suggesting considerable overlap in target loci. Although not directly observable with the structure of our data, the trajectory of some *de novo* mutations is consistent with clonal interference (Gerrish & Lenski, 1998), as observed in other long-term experiments with yeast (Johnson et al. 2021).

### Adaptation to lithium acetate is characterised by fast evolution of functional haploidy

Of the environments investigated, LiAc 0.02M had the most conspicuous changes: four out of five populations were depleted of all standing genetic variation at a remarkably fast pace, likely through a transition to haploidy or fully homozygous diploidy, and always with a *MATα* mating type. The prevalence of the same mating type in this environment may indicate higher fitness of *MATα* (which came from the SK1 parent) or any variant linked to it. Alternatively, the fixing of the *MATα* allele across replicates was a chance event (binomial probability of 0.156).

As none of the LiAc 0.01M populations showed an equivalent response, the fast depletion of diversity in LiAc 0.02M might indicate that this is a dose-dependent response to LiAc. Previous studies have shown that adaptation is typically faster in haploids than in diploids under large population sizes (Johnson et al., 2021; Zeyl et al., 2003). This is because mutations are readily exposed to selection in haploids regardless of their dominance coefficient, and there are multiple other ploidy-dependent effects that can make haploidy advantageous, such as protein-dosage changes and cell size (discussed by Gerstein et al., 2011). Although we did not confirm the ploidy of populations in our experiment, the loci involved were clearly not as advantageous in a heterozygous state. We note, however, that the fitness of the one population (replicate 3) that remained diploid, was just as high as that of the other populations, suggesting that functional haploidy is not the only route for adaptation in LiAc 0.02M. Haplotype trajectories in the replicate 3 population are consistent with the fixation of a single diploid genotype, with an overall loss of 75% of the standing genetic variation (**Figures 3** and **4**). Ultimately, dissecting the genetic basis of the response to LiAc will be necessary to explain these patterns.

### Parallelism at the phenotypic and genomic level

At the phenotypic level, we observed similar fitness responses within environments, especially at the earliest time point measured (generation 100), and most conspicuous in the high-stress environment NaCl and LiAc 0.02M (**Figure 2A**). This suggests that phenotypic parallelism was primarily driven by standing variation. After this, replicate populations diverged significantly in fitness (**Figure 2B-D**). This is interesting because most populations were already fixed for a single genotype at this point, which suggests that it was the influx of *de novo* mutations that diversified phenotypic evolution. In addition, similar fitness values at early stages of evolution might result from genetic redundancy, with multiple subsets of small-effect loci contributing to adaptation but leading to the same phenotype (Barghi et al., 2019). This last scenario might apply to some NaCl populations, where the phenotypic response was highly parallel but multiple genotypes persisted even after 700 generations.

Parallelism at the gene and pathway level is often observed in experimental evolution of *Escherichia coli* and yeast (Huang et al., 2018; Kryazhimskiy et al., 2014; Tenaillon et al., 2012). Here, we detected multiple mutations in genes associated functionally to the HOG pathway, in particular through the Cyc8p-Tup1p and Isw2p-Itc1p complexes. Interestingly, we also observed parallelism at both these levels *across environments*, NaCl and the two concentrations of LiAc, most notably through multiple mutations disturbing the *ISW2* gene. This likely reflects the fact that the stress response system of *S. cerevisiae* is highly pleiotropic. For example, in addition to the stress response, *ISW2* is required for early stages of sporulation (Trachtulcová et al., 2000) and the silencing of subtelomeric genes (Iida & Araki, 2004). Likewise, while the HOG pathway gets activated by osmotic stress, it is also important for the response to heat and oxidative stress, among other stressors (Hayashi & Maeda, 2006; Hohmann, 2009). In an experiment to adapt *S. cerevisiae* to increasing temperature, Huang et al. (2018) detected *de novo* mutations affecting Swip/Snfp, another chromatin remodelling complex heavily involved in multiple stress responses. Swip/Snfp happens to be recruited by the product of *SKO1*, one of the genes mutated in our experiment and repressed by Cyc8p-Tup1p (Proft & Serrano, 1999). Moreover, independent populations of the NaCl environment and in the temperature treatment of Huang et al. (2018) acquired mutations affecting members of the calcium-calcineurin pathway (*CNB1* in our case and *CRZ1* in theirs), once again highlighting the high level of pleiotropy associated with the yeast stress responses.

By contrast, in the EtOH environment we observed mutations affecting a different set of genes implicated in other processes, in particular ubiquitination. It has been shown that ethanol can partially repress the HOG pathway activation (Hayashi & Maeda, 2006), implying that the physiological response necessary for ethanol adaptation might be very different to that of NaCl, LiAc and even heat. Future research could centre on potential trade-offs between adaptation to ethanol and the other environments. In addition to *de novo* point mutations and small INDELs, we also detected cases of parallelism in chromosome copy number variation (**Figure 7**, **Supplementary Figure 8**). Aneuploidies of chromosome 10 in EtOH and chromosome 3 in LiAc 0.01M were maintained for hundreds of generations in two replicates each. Aneuploidies are costly to maintain as they can lead to reduced rate of cell division and cause havoc in gene expression (Dürrbaum & Storchová, 2016; Pavelka, Rancati, & Li, 2010; Pavelka, Rancati, Zhu, et al., 2010; Sheltzer et al., 2011; Torres et al., 2007). However, there is increasing evidence that aneuploidies can facilitate adaptation to environmental stress in yeast (reviewed in Gilchrist & Stelkens, 2019) and drive fitness increases in lab based studies (Chen et al., 2012; Lauer et al., 2018; Millet et al., 2015; Selmecki et al., 2009) and wild populations (Gasch et al., 2016; Scopel et al., 2021; Tsai & Nelliat, 2019). Since aneuploidies are costly, the temporal patterns and parallelism we observe across replicates suggest they had adaptive benefits.

We detected no direct link between the distribution of *de novo* mutations and aneuploidies, suggesting that this parallelism is driven by genetic variation that pre-existed in the founder, and became adaptive through changes in copy number. Similarly, the repeated transition towards functional haploidy in LiAc 0.2M populations was likely facilitated by the ancestral diversity in the founder, as the change was remarkably fast. Overall we see evidence of parallelism at the phenotypic and molecular level derived from both standing genetic variation and *de novo* mutations.

### Interaction between standing genetic variation and de novo mutations

One unexpected aspect of our data is the changing trajectory of some haplotypes, most notably in LiAc 0.01M populations, but also in some NaCl populations (**Figure 3**). We found no evidence that methodological artefacts, namely low depth of coverage, explain this pattern. However, we did observe simultaneous emergence of *de novo* mutations that follow the trajectory of changing haplotypes of standing genetic variation. Just like mutations can compete in otherwise isogenic populations (i.e. clonal interference), these mutations might have changed the trajectory of resident haplotypes to the detriment of some others, thus contributing to the maintenance of variation for longer periods.

Some of the unexpected changes in haplotype frequencies may also be explained by fluctuating selection caused by the evolving populations themselves, for instance through changes in cell metabolism impacting the pH of the growth medium, which may have altered the direction of selection for some haplotypes. Changes in the growth medium may affect yeast populations in similar ways as seasonal changes cause trait and genomic shifts in populations of fruit flies as recently shown in an elegant field experiment, where the continuous adaptation in response to rapid environmental change was described as adaptive tracking (Rudman et al., 2022).

## Conclusion

Our study shows that adaptation dynamics are affected by an intricate interplay of standing genetic variation, *de novo* mutations, and different levels of environmental stress. Follow up studies are needed to provide a better understanding of the relative roles that each of these factors, and their interactions, play in changing a population’s adaptive potential to future environmental change. Since we found considerable variation in population responses to different concentrations of one stressor (lithium acetate), one interesting angle would be to use an evolve-and-resequence approach along a fine-scaled environmental gradient of stress, to mimic population responses to gradual pollution events and environmental deterioration. Similarly, one could create a gradient of standing genetic variation, ranging from clonal to increasingly outbred populations, to conclusively test how increasing genetic diversity affects adaptation dynamics. Pairing experimental evolution with long read sequencing could be used to scrutinise the role of larger structural variants beyond small INDELs in adaptation dynamics (for instance transposable elements, inversions and translocations), to conclusively test for their causality in fitness changes and adaptive processes. We hope our study contributes to a better understanding of the constraints and drivers shaping the evolutionary potential of populations and their responses to future environmental change.

## Acknowledgements

We thank Duncan Greig for providing the SK1 and Y55 strains. We thank Maria de la Paz Celorio Mancera, Viktoria Köppä, and Zebin Zhang for helping with serial transfers and media preparation. We thank Dmitri Petrov, Anthony Long, Noah Gettle and Milo Johnson for valuable discussion of the genomic data. Computation was performed on resources provided by SNIC through Uppsala Multidisciplinary Center for Advanced Computational Science (UPPMAX) under Projects snic2019-30-49 (Small Storage), snic2021-6-267 (Medium Storage), snic2019-8-355 and snic2022-22-46 (Computation). We acknowledge support from Science for Life Laboratory, the National Genomics Infrastructure, NGI, and Uppmax for providing assistance in massive parallel sequencing and computational infrastructure.

## Data statement

Phenotypic data are available in **Supplementary Table 1**. The raw sequencing data were deposited in the European Nucleotide Archive, accession number PRJEB46680. The code used for alignment, variant calling, analyses and plotting are available at https://github.com/SLAment/AdaptationDynamics.

## Author Contributions

RS and CG conceived and designed the study. CB constructed the founder populations. CG carried out the serial transfers with help from CB, JSG, and RS. CG collected and analysed the fitness data with help from JGS. CG and JGS extracted DNA. SLAV and NR carried out alignments, variant calling and filtering. SLAV and AR analysed genomic dynamics. DPB performed aneuploidy analyses. CG, SLAV and RS wrote the manuscript and visualised the results with help from all authors. CG is responsible for the data curation. RS acquired the funding and resources for this study and provided supervision. All authors gave final approval for publication and agreed to be held accountable for the work performed therein.

## Supplementary Figures

**Supplementary Figure 1.**
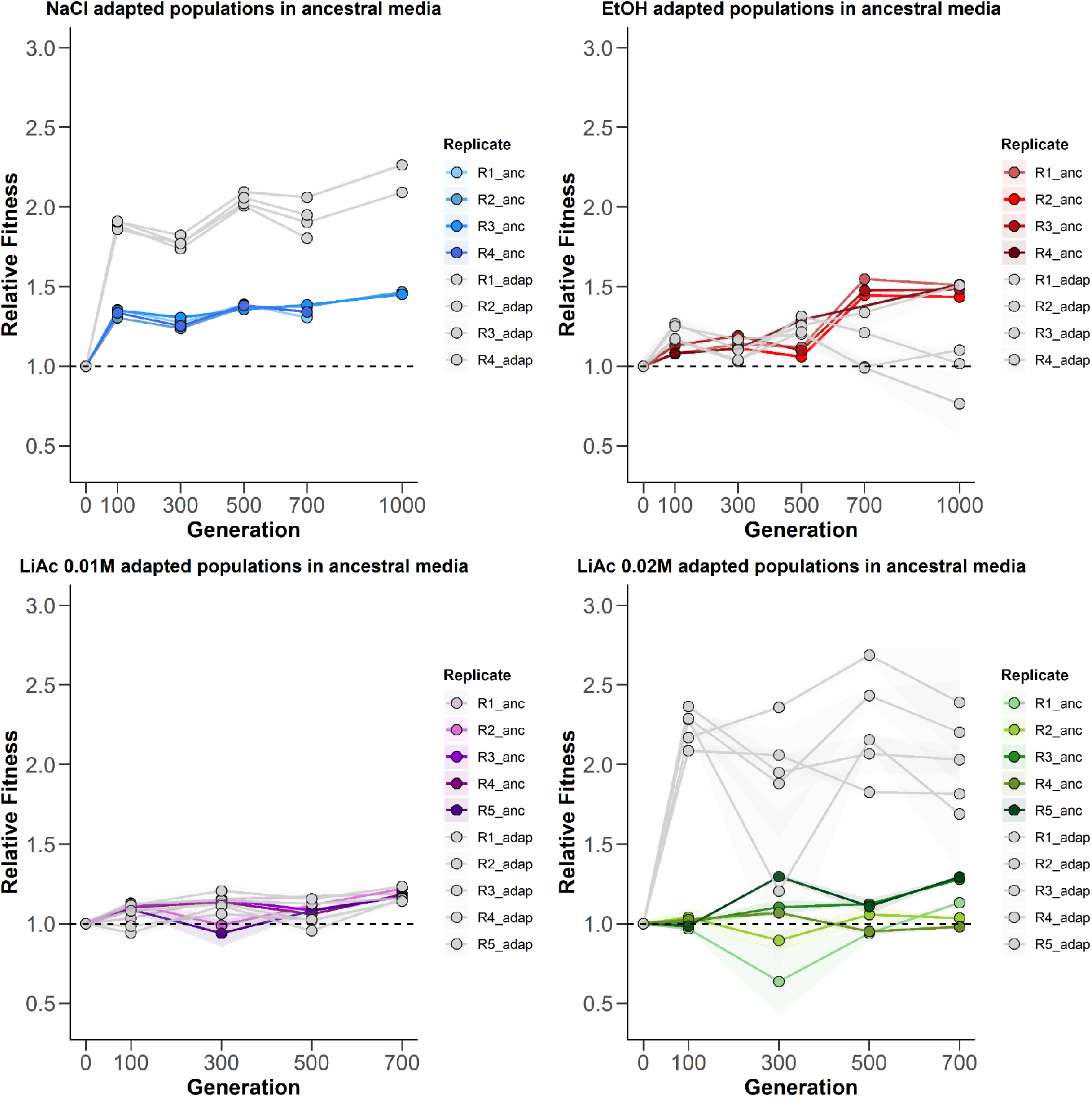
Mean relative fitness (optical density after 24h vs. founder populations) of evolved populations in ancestral (SC media) and selective conditions. Evolved populations tested in ancestral conditions are shown in different shades of colour (NaCl = blue, EtOH = red, LiAc 0.01M = purple, LiAc 0.02M = green). Evolved populations tested in selective environments are shown in grey (cf. Figure 2). Shaded areas are 95% confidence intervals. One EtOH population went extinct in the first 100 generations. Two NaCl populations after 700 generations were removed due to contamination.

**Supplementary Figure 2.**
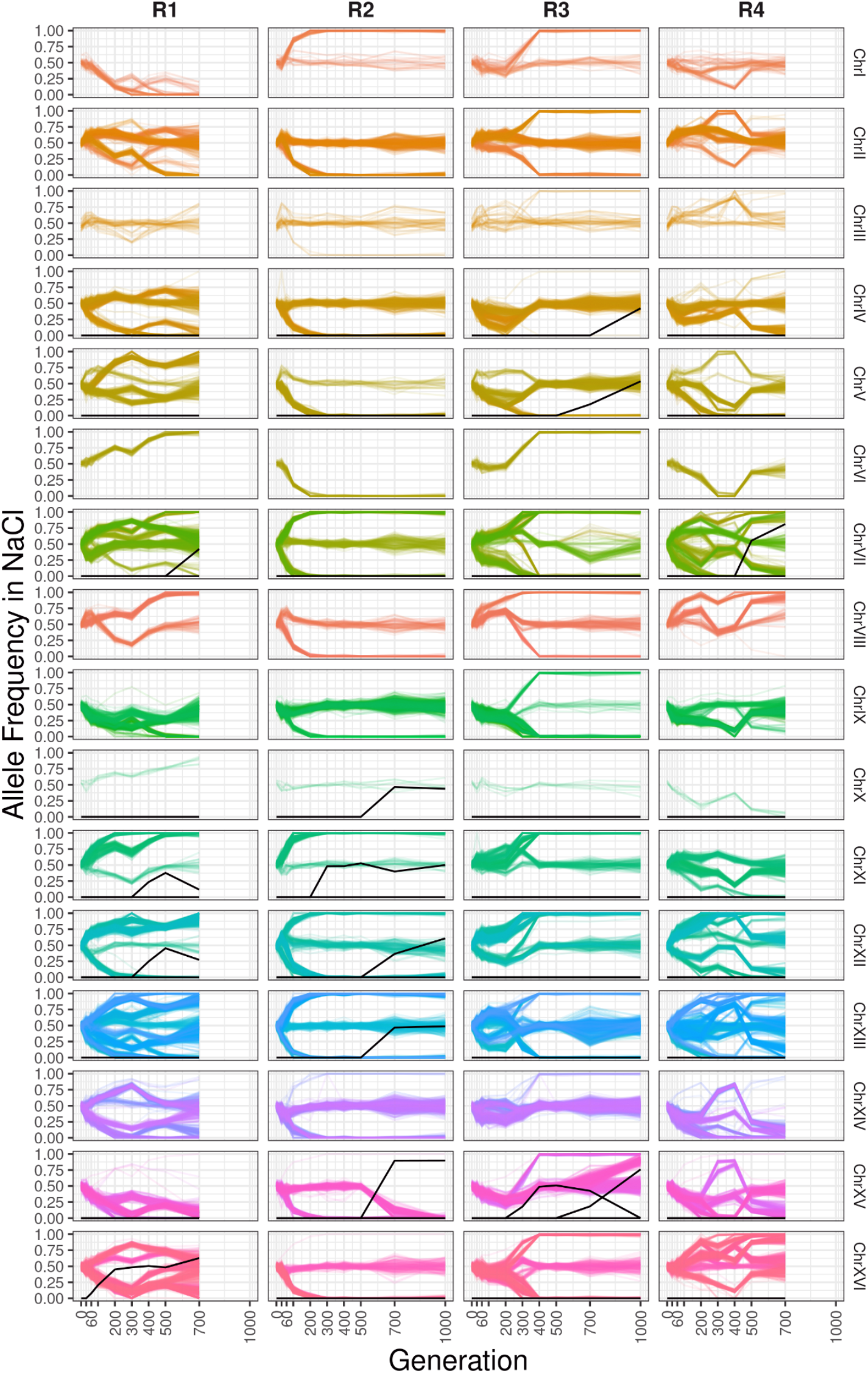
Allele frequency trajectories from the ancestral SNP variation (colours) and *de novo* mutations (MAF >0.35, black) in the NaCl environment. Lines connect the allele frequencies of all sites per chromosome and per replicate. Only the allele of the parental strain SK1 is shown in the case of ancestral variation.

**Supplementary Figure 3.**
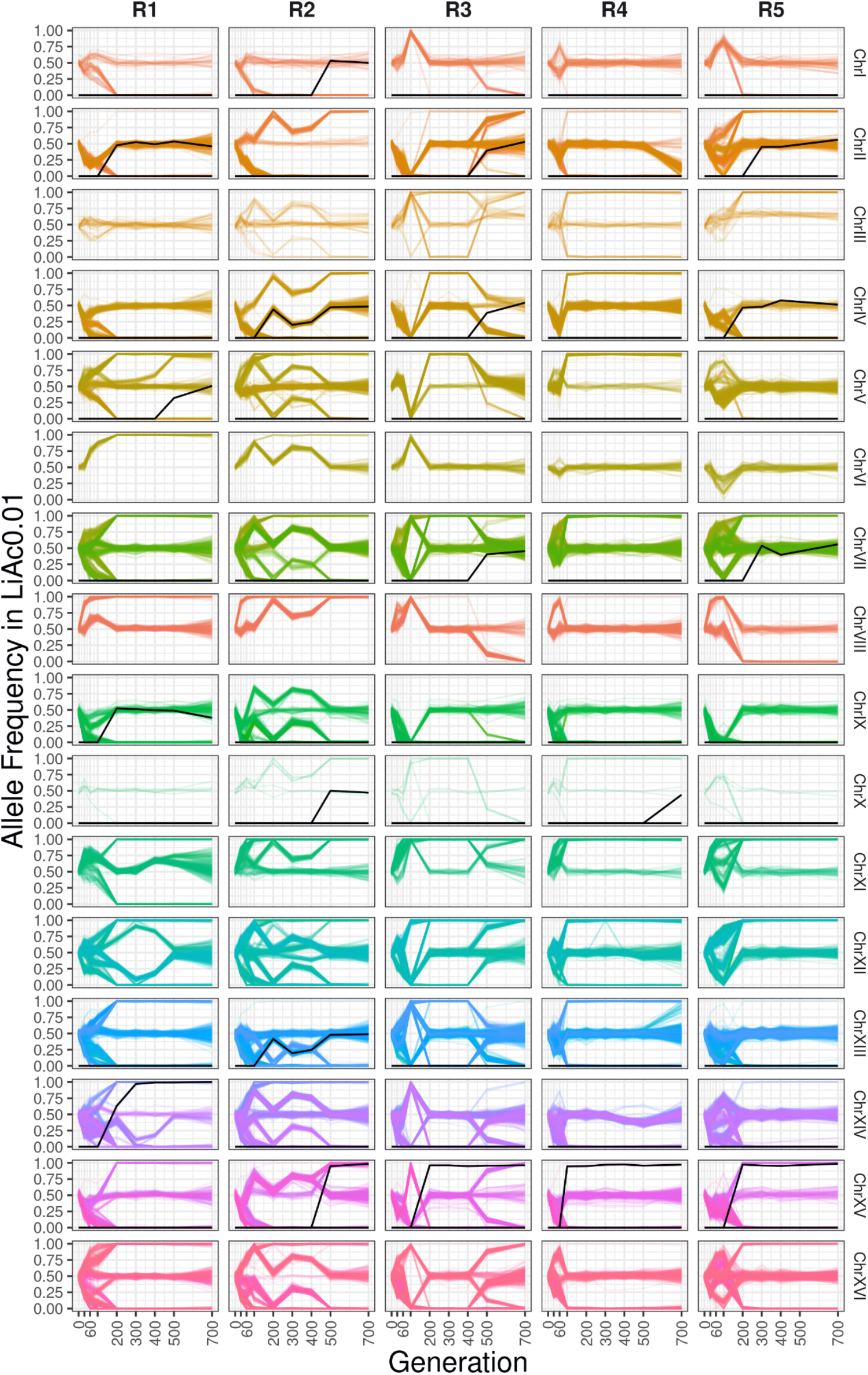
Allele frequency trajectories from the ancestral SNP variation (colours) and *de novo* mutations (MAF >0.35, black) in the LiAc 0.01M environment. Lines connect the allele frequencies of all sites per chromosome and per replicate. Only the allele of the parental strain SK1 is shown in the case of ancestral variation.

**Supplementary Figure 4.**
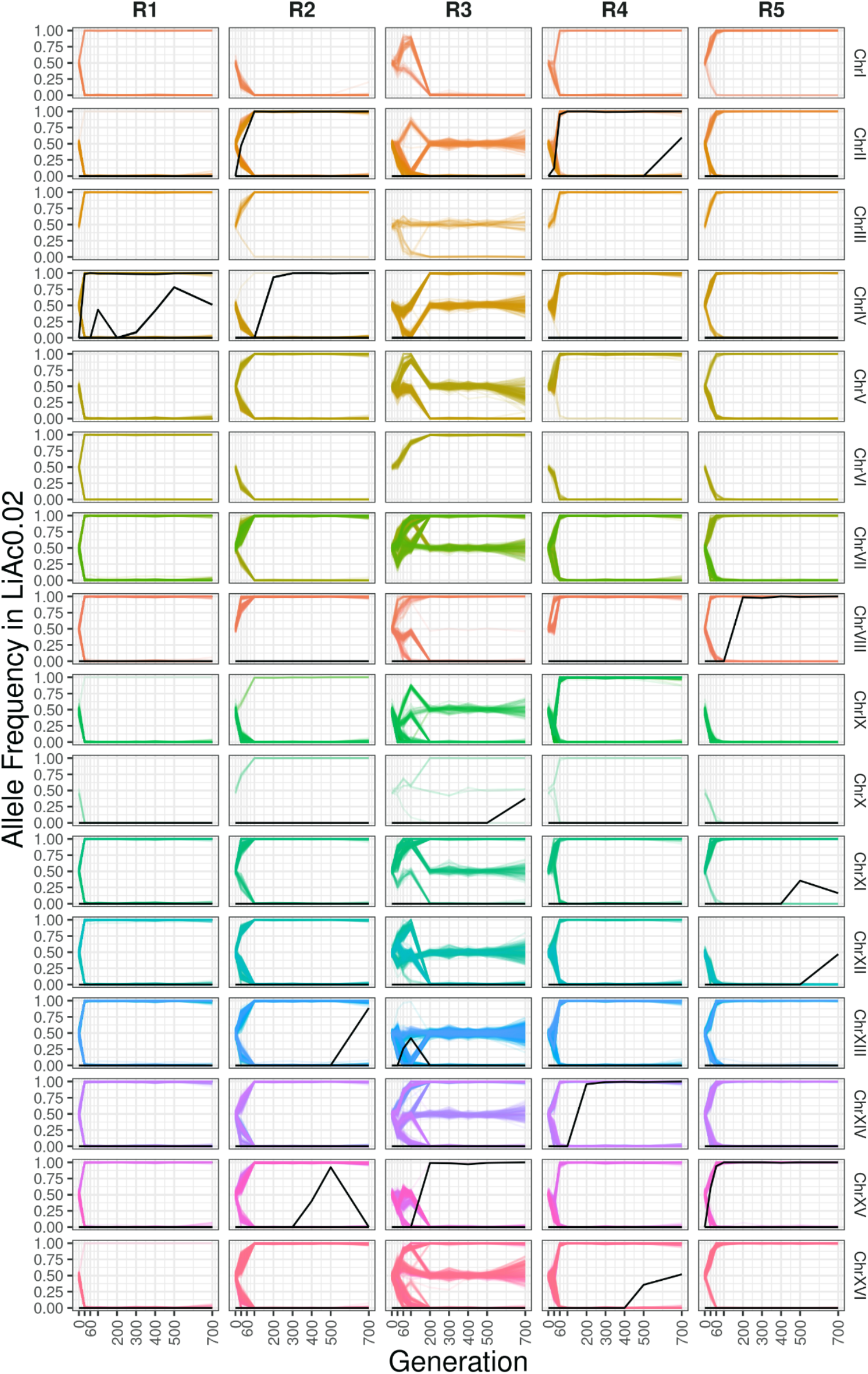
Allele frequency trajectories from the ancestral SNP variation (colours) and *de novo* mutations (MAF >0.35, black) in the LiAc 0.02M environment. Lines connect the allele frequencies of all sites per chromosome and per replicate. Only the allele of the parental strain SK1 is shown in the case of ancestral variation.

**Supplementary Figure 5.**
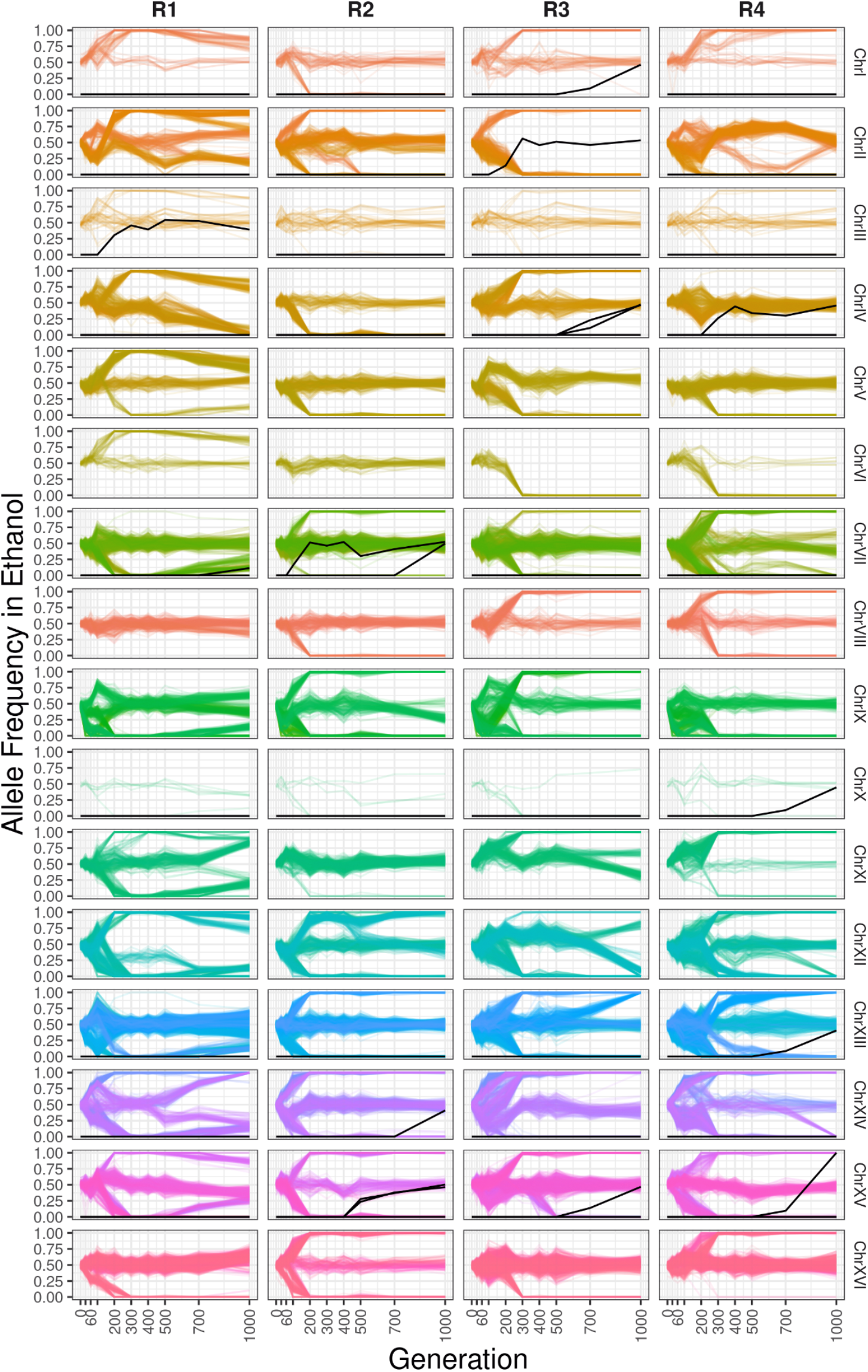
Allele frequency trajectories from the ancestral SNP variation (colours) and *de novo* mutations (MAF >0.35, black) in the EtOH environment. Lines connect the allele frequencies of all sites per chromosome and per replicate. Only the allele of the parental strain SK1 is shown in the case of ancestral variation.

**Supplementary Figure 6.**
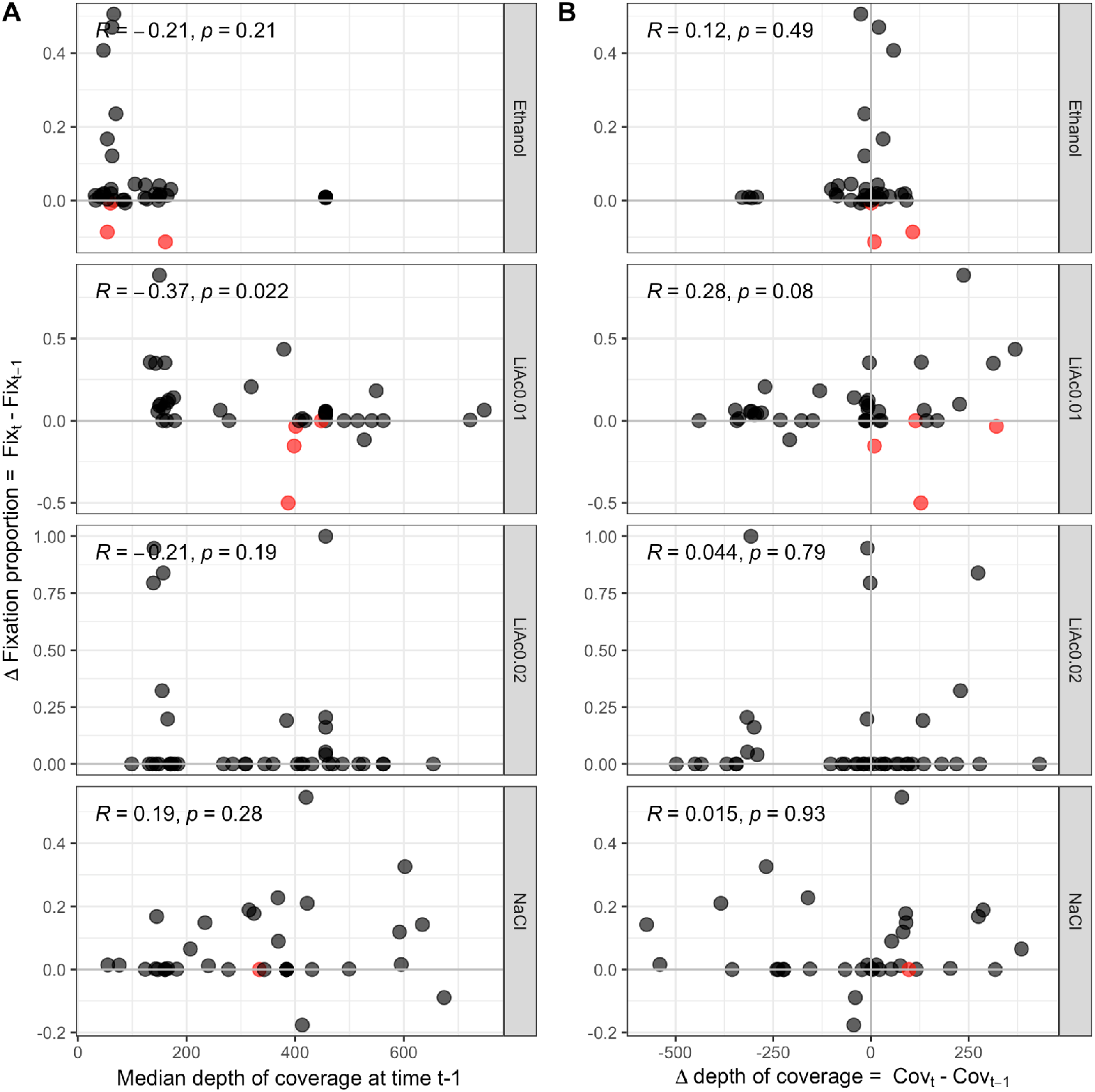
Changes in the proportion of nearly fixed (MAF < 0.1) sites is not explained by the depth of coverage of a sample. For each sample, we calculated the difference between the proportion of nearly fixed sites at time *t* and the proportion at the previous time *t* -1 (*ΔFixation proportion*). (A) If low coverage is the cause of “unfixing” sites (*ΔFixation proportion* < 0), then we should observe a positive correlation between *ΔFixation proportion* and depth of coverage at *t* -1, but this does not seem to be the case. (B) We further calculated the difference in median depth coverage between time *t* and *t* -1 and plotted the relationship with *ΔFixation proportion*. Few samples fall into the lower right corner of individual panels (red points), which once more indicates that negative differences in fixation proportion are not generally associated with negative differences in coverage. Significance of *p-values* correspond to Pearson correlations.

**Supplementary Figure 7.**
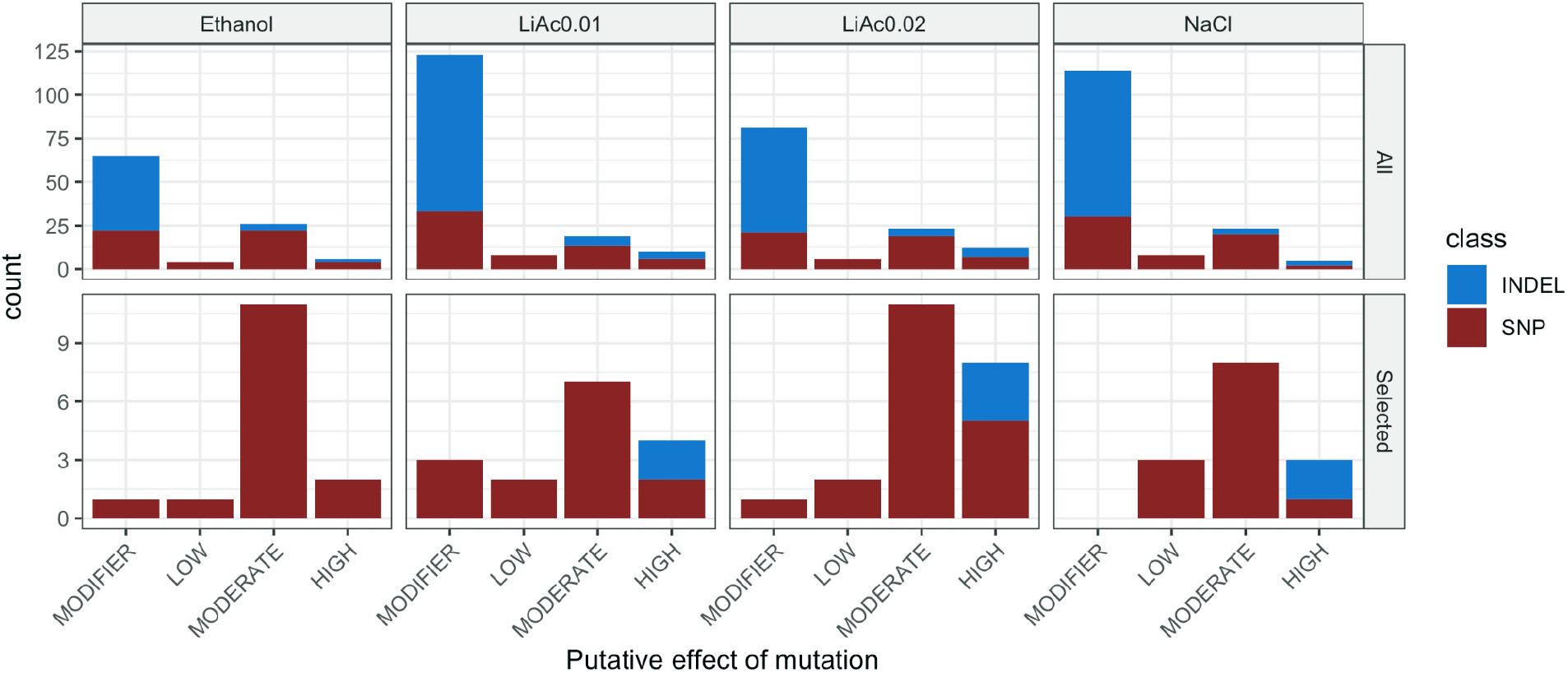
Putative fitness effects of de novo mutations as predicted by SnpEff. The upper panel shows the effect of all mutations identified, while the bottom includes only those that reached a frequency of at least 35% during some point of the experiment and that passed the manual curation. “Modifier” indicates mutations falling in the non-coding region of the gene (e.g. enhancers/repressors), “Low” is synonymous mutations, “Moderate” includes missense mutations and in-frame deletions/insertions, and “High” is frameshift variants and gain or loss of stop codons.

**Supplementary Figure 8.**
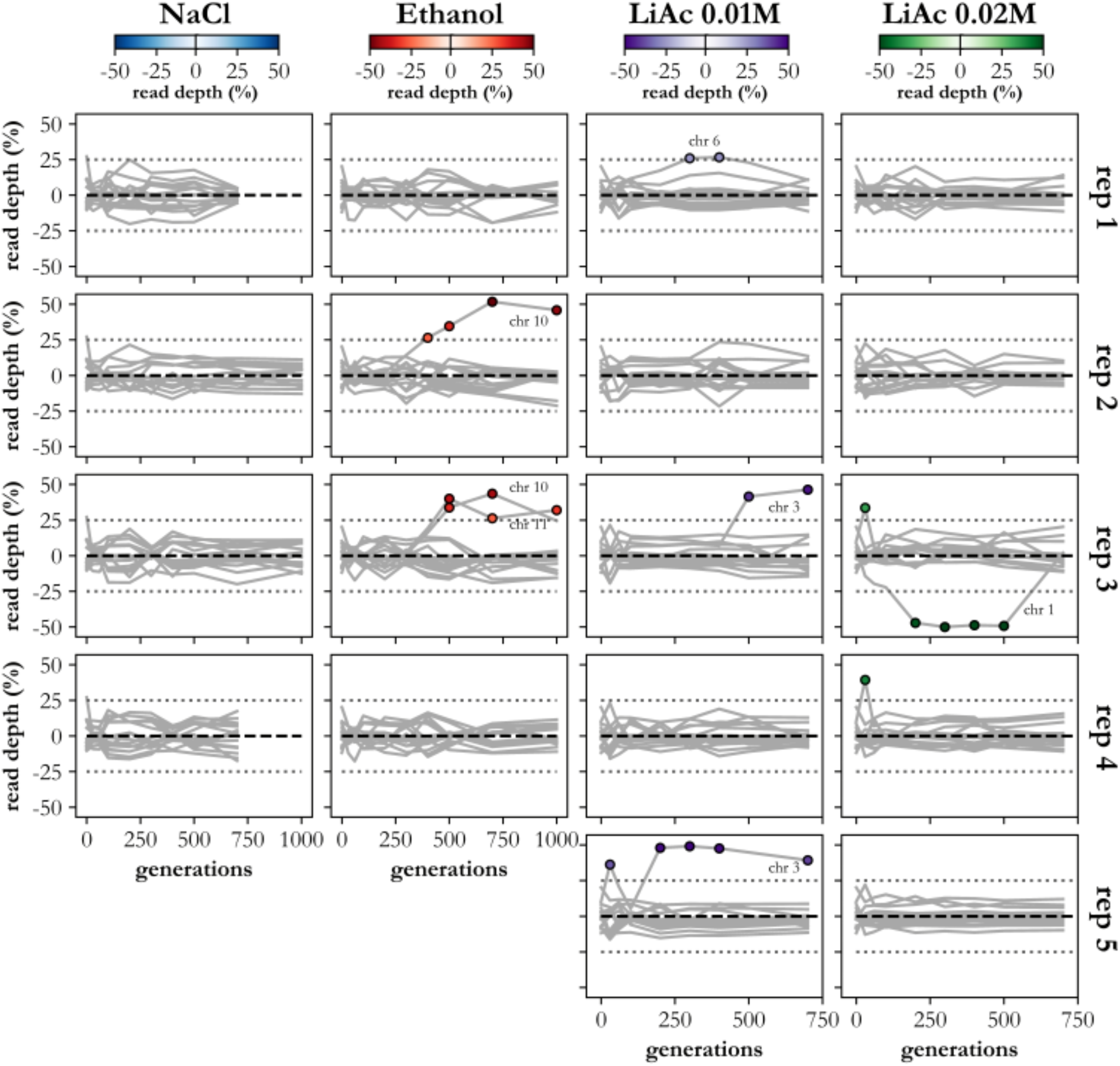
Relative chromosomal read depth during adaptation. Lines show the relative read depth of each chromosome (read depth vs. the mean read depth across all chromosomes). Chromosomal deviations greater than 25% from genome read depth are indicated with circles and colour shade indicates the severity of deviation.

**Supplementary Table 2.**
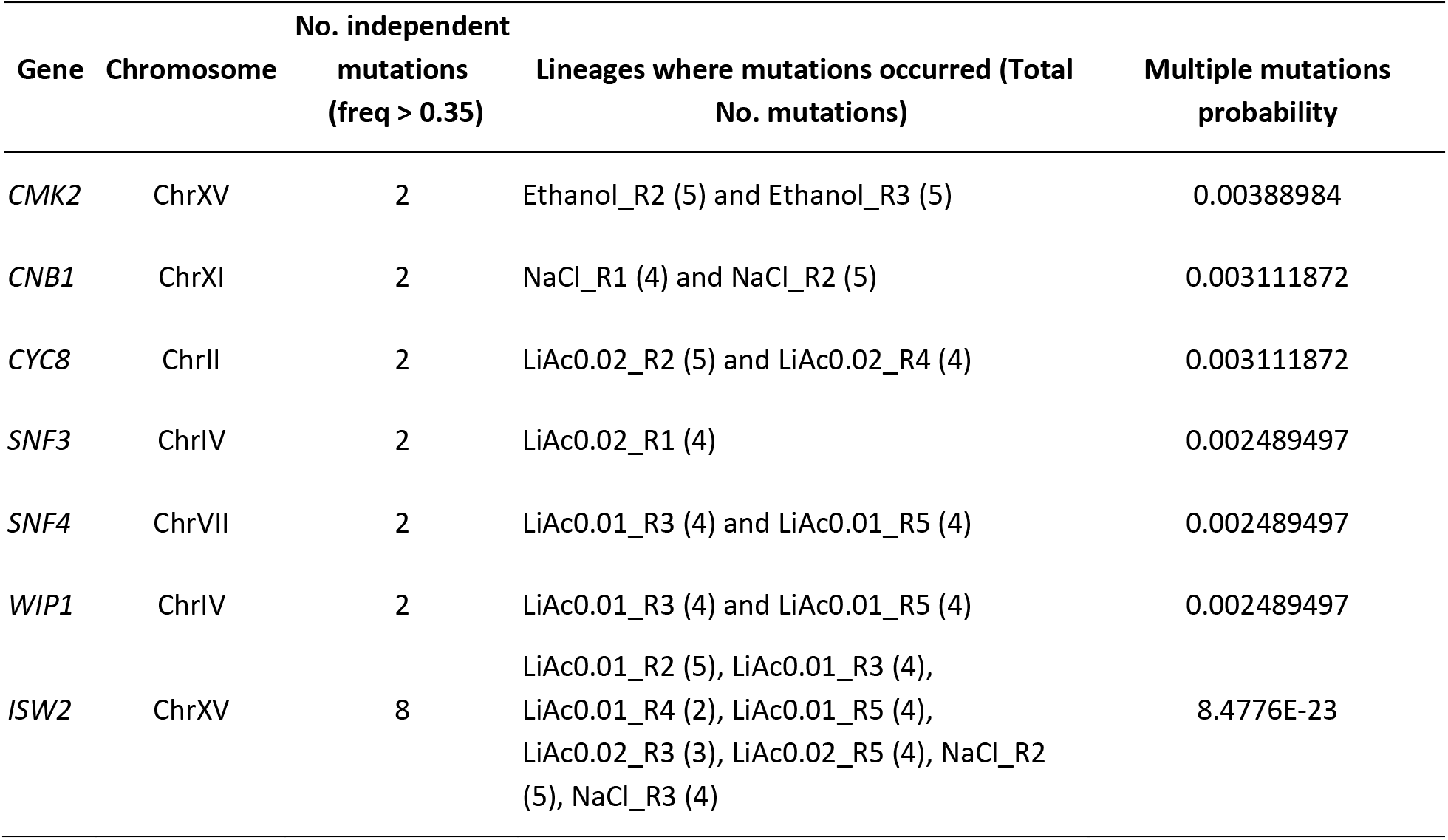
Genes with independent high frequency (>0.35) de novo mutations and associated probability of independent mutations per gene.

